# A Relational Code for Objects in Primate Frontal Cortex

**DOI:** 10.64898/2026.10.17.759839

**Authors:** Nicholas Watters, Jack Gabel, Joshua Tenenbaum, Mehrdad Jazayeri

**Affiliations:** Department of Brain and Cognitive Sciences, Massachusetts Institute of Technology, Cambridge, MA, USA; McGovern Institute for Brain Research, Massachusetts Institute of Technology, Cambridge, MA, USA; Howard Hughes Medical Institute, Massachusetts Institute of Technology, Cambridge, MA, USA

**Author notes:** Correspondence (MJ).

## Abstract

When we remember a scene, we recall both the locations of individual objects and their spatial arrangement. Yet, how neurons encode this arrangement alongside individual locations is unknown. In this work, we recorded large-scale neural activity from the frontal and supplementary eye fields of monkeys performing a two-object spatial working-memory task. Using a continuum of two object configurations, we tested whether neurons encoded individual object locations or the spatial relationship between them. Many neurons had well-defined position tuning when a single object was presented. With two objects, responses spanned a continuum from retinotopic to relational tuning. Surprisingly, most units were better explained by relational tuning than retinotopic tuning. Relational and retinotopic units encoded complementary dimensions of the stimulus, and their observed mixture supported near-optimal population decoding of object locations. Relational tuning was systematically aligned with single-object receptive fields, enabling a readout that produced repulsive biases resembling those observed in behavior. Standard multi-object models such as averaging and normalization failed to reproduce the observed continuum, whereas a model integrating receptive fields over potential saccades captured it. These findings show that the frontal oculomotor cortex represents not only where objects are, but also how they are arranged with respect to one another.

## INTRODUCTION

A glance at a desk reveals a scene composed of multiple objects: a mug beside a laptop, a lamp behind them, a notebook to the side. How is the spatial information in such a scene organized in the mind? The brain might represent a collection of individual object locations in a common reference frame, such as gaze direction. Or it might represent the objects relative to one another, e.g. the mug is to the left of the laptop. The first format preserves the independent locations of objects; the second captures their spatial relations and remains invariant to changes of the observer’s reference frame. These formats are complementary, yet little is known about whether, and where, the primate brain represents spatial relations between objects.

The importance of object-relative coding was apparent in early work on dorsal-stream function. Ungerleider and Mishkin found that posterior parietal lesions impaired monkeys’ ability to choose a target specified relative to a landmark^1^. Subsequent physiology, however, focused primarily on how the position of an individual object is represented in a common eye or head coordinate system. Variations in receptive fields were interpreted as transformations between coordinate systems or remappings that update position codes^2,3^. These interpretations retained the central premise that object locations are represented within observer-centered frames while setting aside whether neurons represent the positions of objects with respect to one another.

The frontal eye field (FEF) and supplementary eye field (SEF) are natural substrates to revisit this question. Both areas contain neurons with spatially selective visual, memory, and saccade-related responses, tuned in eye-centered or head-centered coordinates^4–10^. However, their spatial selectivity is not always described by a stable observer-centered receptive field. FEF receptive fields are updated around saccades^11–13^, and visual landmarks influence target selectivity in both FEF and SEF^14–16^. Moreover, some SEF neurons are modulated by whether a target lies to the left or right of a reference object despite changes in its retinotopic position^17–19^. While these results suggest relational coding is plausible, whether FEF or SEF systematically code for vectors between objects is not known.

Detecting such a code requires a task with multiple behaviorally relevant objects whose absolute and relative positions are systematically varied. Here, we performed large-scale single-unit recordings from monkeys’ FEF and SEF in a spatial working-memory task with two objects whose positions varied parametrically, both in absolute coordinates and relative to one another. This design allowed us to distinguish responses predicted by the retinotopic positions of the objects from those predicted by the vector between them.

We found that relational tuning was prevalent in both areas and that responses spanned a continuum from retinotopic to relational. Relational and retinotopic units encoded complementary dimensions of the stimulus, and their observed mixture supported near-optimal population decoding of stimulus locations. Moreover, alignment between relational tuning and single-object receptive fields predicted a repulsive bias in remembered locations, paralleling a bias in the animals’ behavior. Standard implementations of averaging, normalization and inhibitory receptive-field models could not account for relational coding. By contrast, models in which retinotopic fields were modulated by inter-object orientation, or by the expected sensory consequences of planned saccades, captured the observed relational responses. Together, these findings show that, in a multi-object working-memory task, the frontal oculomotor cortex represents not only where individual objects are, but also how they are arranged with respect to one another.

## RESULTS

### Task and Behavior

We trained two rhesus macaques on a multi-object spatial working-memory task (Fig. 1a). Each trial began with central fixation, followed by a 1-sec stimulus period in which one or two objects were presented at locations randomly sampled from a ring (Fig. 1b). The objects then disappeared for a 1-s delay, after which, a central cue identified one object as the target. Monkeys had to make a saccade to that target’s remembered location to receive a juice reward. On two-object trials, either object could be cued as the target; thus animals had to remember the location of both objects. Because gaze was held at fixation throughout the stimulus and delay periods, retinotopic and craniotopic coordinates were indistinguishable; we therefore refer to all absolute spatial coordinates as retinotopic.

**Figure 1.**
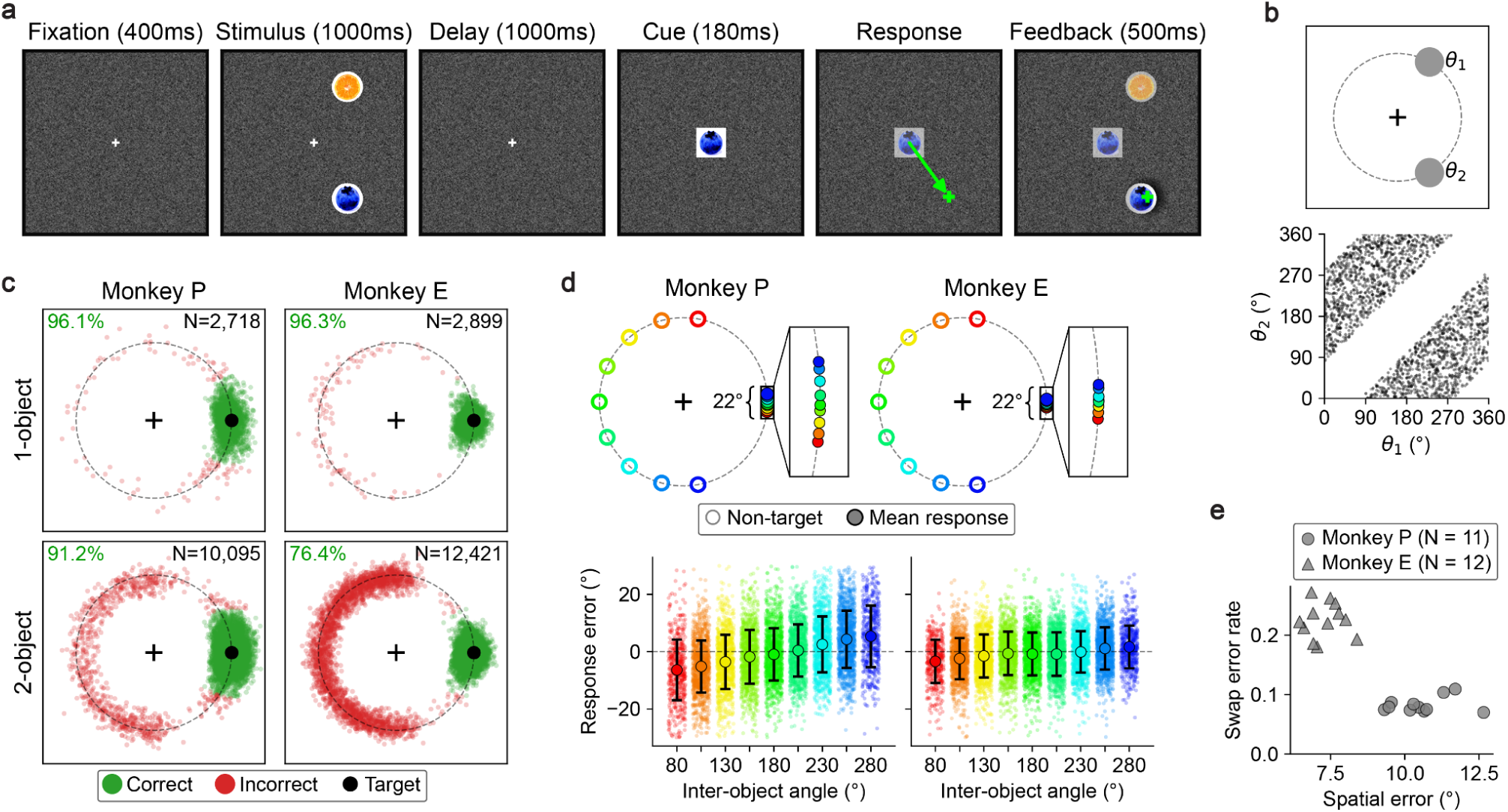
Task and behavior. **(a)** Example trial (left to right). Gaze fixation must be held near the central cross from trial onset until response. The green arrow in the response phase depicts a saccade vector and was not displayed to the monkey. The green crosses in response and feedback phases show gaze position and were not displayed to the monkey. After the response saccade, all stimulus objects were revealed for visual feedback and a reward was delivered if the response was correct (near the stimulus object matching the cue). **(b)** 2-object stimulus conditions. (top) Dashed circle shows ring from which all object positions were sampled. Gray circles are object positions for an example trial. (bottom) Each point represents a 2-object trial, showing the angular positions of the two objects. **(c)** Responses aligned to target. Each small dot is a response for one trial, rotated around fixation so the cued (target) object lies on the black dot. (top) 1-object conditions for each monkey. (bottom) 2-object conditions. In all plots, the total number of trials is shown in the top-right and the percentage of correct trials is shown in the top-left. **(d)** Repulsion bias. Same data as in (c, bottom), except binned by and conditioned on the angle of the non-target object, excluding trials with angular response error greater than 30° with respect to the fixation cross. (top) Open circles show non-target position and solid dots show mean response angle for each bin. Inset enlarges the region around the target position. (bottom) Each small dot is the response angle relative to the target on one trial, with x-position randomly jittered after binning. Large dots and errorbars show means and standard deviations. Bin mean is −6.4°/-3.4° at 80° and +5.3°/+1.6° at 280° for Monkey P/E. Average repulsion is +4.66° ± 0.18 for Monkey P and +1.50° ± 0.14 for Monkey E. **(e)** Types of response errors for 2-object trials. Each dot reflects one session. Spatial error (x-axis) is the estimated standard deviation of angular responses around the intended response object. Swap error (y-axis) is the estimated probability of responding toward the non-target object (Methods).

Both monkeys performed the task reliably in one- and two-object conditions (Fig. 1c). On two-object trials, saccades to the cued object were systematically biased away from the uncued object, with the largest repulsion when the objects were closest together (Fig. 1d). This pattern parallels repulsive biases reported in human multi-object working-memory tasks^20–23^. Although the two animals showed qualitatively similar repulsion, they differed in their trade-off between spatial error around the cued object and swap errors toward the uncued object (Fig. 1e).

### Neurophysiology

We used Neuropixels probes^24^ and Plexon V-Probes to record large-scale spiking activity simultaneously from FEF and SEF while monkeys performed the task. Some units behaved as expected from a retinotopic receptive field: They responded when one of the objects occupied a preferred location in visual space (Fig. 2a). Other units showed a strikingly different pattern: They responded strongly to object pairs spanning a particular orientation, even when the two objects occupied different retinotopic locations (Fig. 2b). Thus, their preferred spatial configuration was not a location on the display, but the orientation of the line connecting the objects. We refer to this response pattern as relational tuning and to neurons that exhibit it as relational units. Relational units were modulated during both the stimulus and delay periods and were observed in both FEF and SEF (Fig. 2b-d).

**Figure 2.**
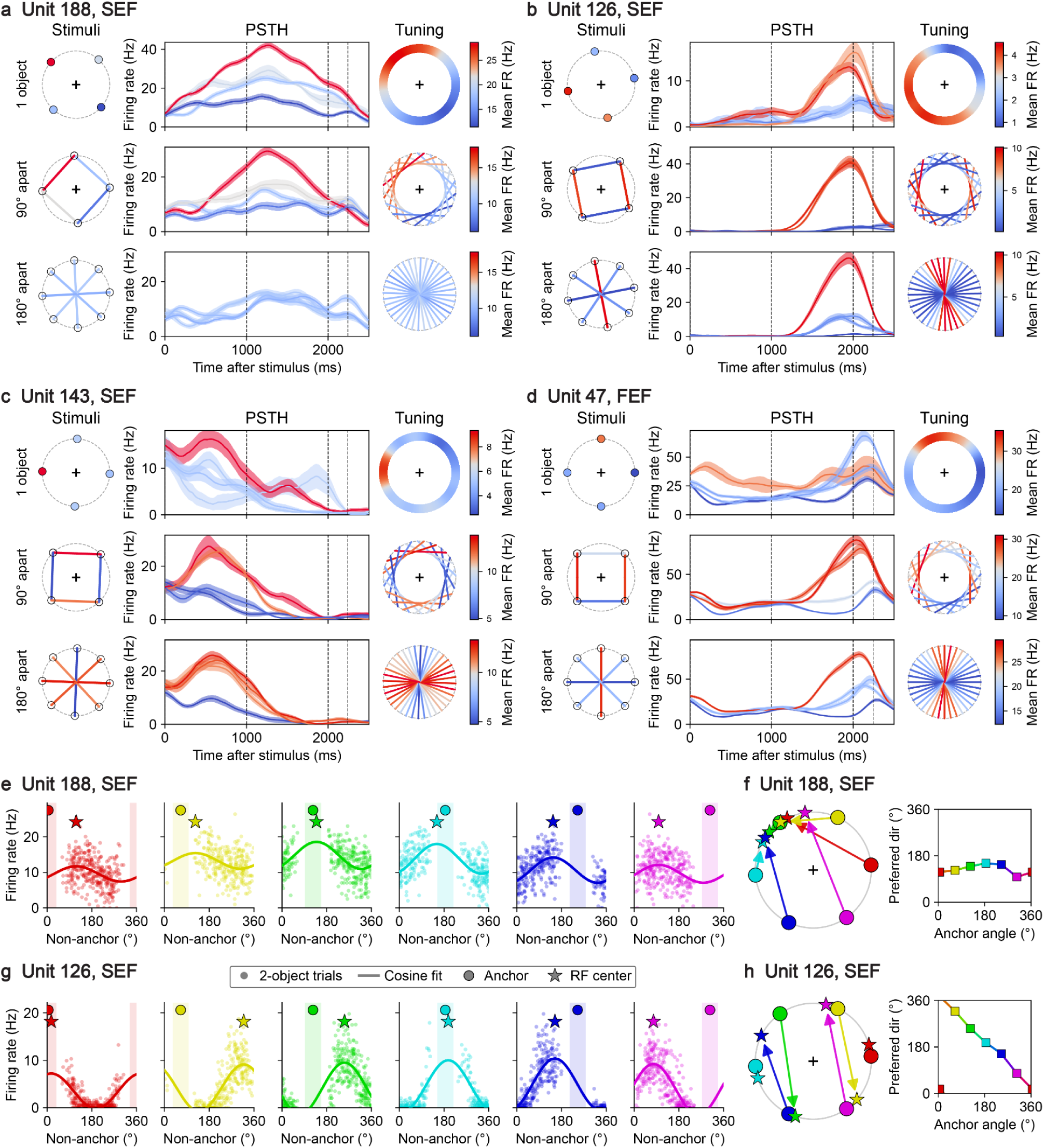
Single units are tuned to relative object positions. **(a)** Unit 188, recorded from area SEF. Rows are families of stimulus conditions: One object (top), two objects 90° apart (middle), and two objects 180° apart (bottom). Left column shows four example stimuli. 2-object stimuli are represented by a chord between the two objects. Colors show mean firing rate over stimulus and delay phases for trials where the objects are within 15° of the shown stimulus. Middle column shows peri-stimulus time histograms (PSTHs) for the conditions in the left column, smoothed with a 250ms triangular kernel. Solid lines show means, and errorbands show 95% confidence intervals of those means. Dashed vertical lines indicate delay onset time, cue onset time, and mean response time. Right column is analogous to the left column except for more stimuli. For the 1-object condition, 180 stimuli are shown. For the 2-object conditions, 18 stimuli are shown. All are colored by mean firing rate. **(b-d)** Analogous to panel (a) except for unit 126 (from area SEF), unit 143 (from area SEF), and unit 47 (from area FEF), respectively. **(e)** Conditional 2-object responses for unit 188. Each plot corresponds to a different anchor point (large dot) and shows 2-object trials where one object lies near that anchor (within the shaded region). Each small dot shows the angle of the other object (x-axis) and mean firing rate (y-axis) for each trial. Curved line shows a cosine tuning curve fit to the small dots, with maximum indicated by the star. **(f)** (left) Each arrow points from an anchor (dot) to the corresponding conditional tuning curve peak (star). (right) Same data and color scheme as left, except shown in a lineplot format. Preferred direction (y-axis) is the angle of the conditional tuning curve peak. **(g)** Analogous to panel (e) except for unit 126. **(h)** Analogous to panel (f) except for unit 126.

The two coding schemes make contrasting predictions for a unit’s conditional tuning: the preferred location of one object when the position of the other object is specified. If a unit is retinotopic, this conditional preferred direction should remain fixed. If a unit is relational, this conditional preferred direction should instead move around the ring in the opposite direction of the conditioned object, thereby preserving the preference for the orientation of the line connecting the objects. The observed responses followed these predictions: Retinotopic units maintained a constant conditional preferred direction (Fig. 2e,f), whereas relational units showed the predicted counter-rotation (Fig. 2g,h).

### Organization and functional implications of relational coding

#### Prevalence of relational orientation coding

To assess the prevalence of relational tuning in FEF and SEF, we compared three models of responses to two-object stimuli (Fig. 3a): (1) a non-relational retinotopic model for units with a preferred object location, (2) a relational model for units tuned to the orientation of the line connecting the objects, regardless of their locations, and (3) an alternative relational model, termed “proximity,” for units tuned to the distance between the objects, regardless of their locations or the orientation of the connecting line.

**Figure 3.**
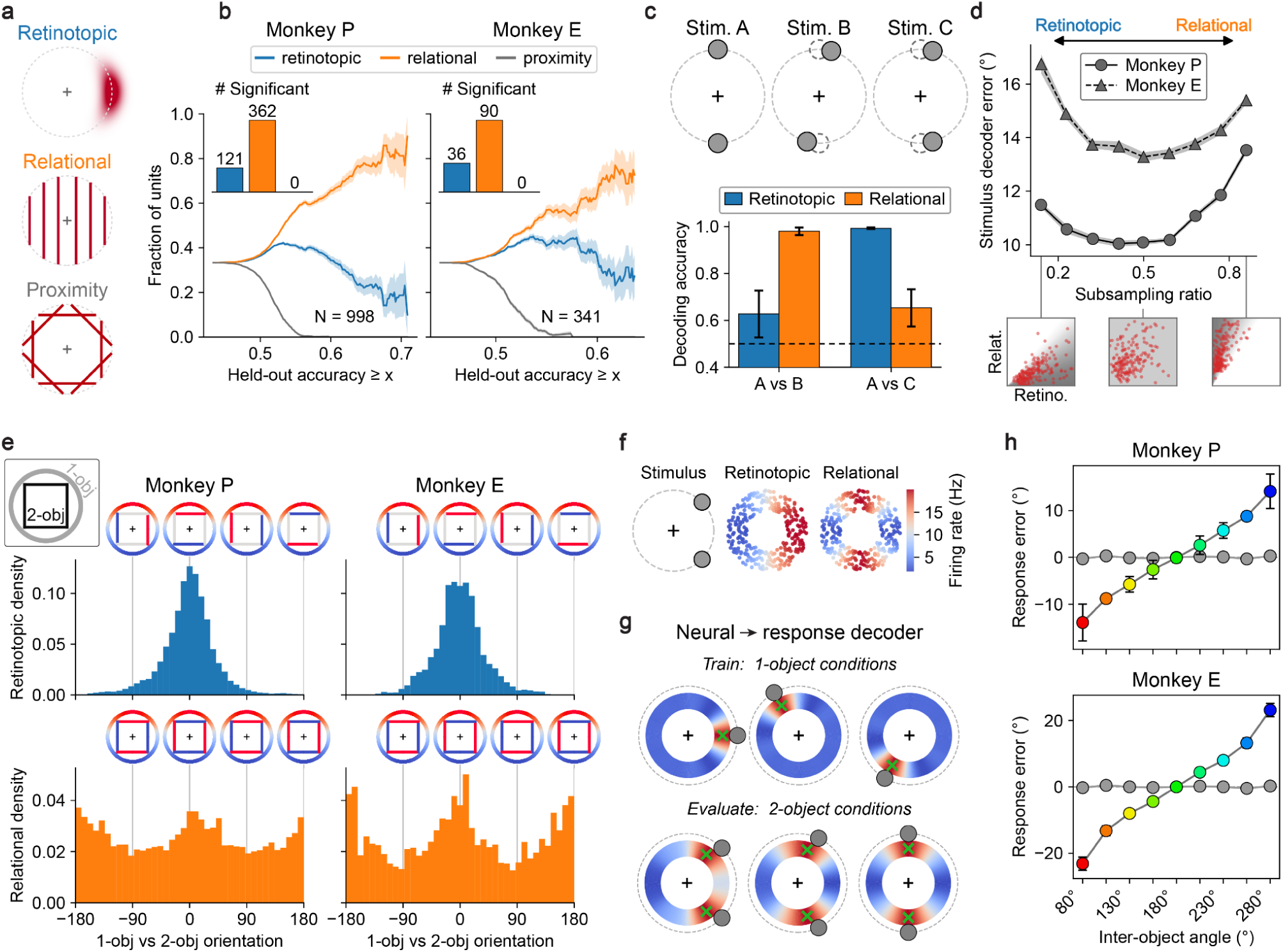
Population activity. **(a)** Retinotopic (top), relational (middle), and proximity (bottom) models. Each plot shows an example idealized unit. In the retinotopic model, the shaded region depicts a receptive field. In the relational and proximity models, chords show preferred 2-object stimuli. **(b)** Prevalence of relational units for Monkey P (left) and Monkey E (right). X-axis is a lower bound on cross-validated model accuracy (chance is 0.5). Each x-value determines a subset of (unit, model) pairs. The y-value shows the fraction of the subset corresponding to each model. Errorbands show s.e.m. Inset shows the number of units for which each model had significantly higher (p < 0.05) cross-validated accuracy than both other models. **(c)** What each code distinguishes. (top) Three example 2-object stimuli. In stimulus B and C, the dashed circles show the object positions in stimulus A. (bottom) Cross-validated accuracy of a linear decoder trained to discriminate pairs of stimuli from the responses of the top 10 retinotopic neurons from each monkey (blue) and top 10 relational neurons from each monkey (orange). Errorbars show standard error of the mean. **(d)** Relational units enable efficient stimulus representation. Decoder error (y-axis) is the bootstrapped mean angular error per object of an MLP trained to decode 2-object stimuli from a fixed-size sub-population of units. The subsampling ratio of the sub-populations (x-axis) is the mean rank of the ratio of relational model accuracy to retinotopic model accuracy of the sampled units. By construction, subsampling ratio 0.5 reflects an unbiased sample of the neural population. Insets below show this sampling distribution (gray shading) and a sub-population draw (red dots) for three sweep points, in the space of retinotopic accuracy (x-axis) versus relational accuracy (y-axis). **(e)** Alignment between 2-object and 1-object receptive fields. (inset) Square represents four distinct 2-object stimuli, circle represents 1-object tuning, as in Figure 2a-d. (top) Receptive field alignment between 1-object conditions and 2-object conditions for units where the retinotopic model significantly outperforms a constant null (N=623 for Monkey P and 153 for Monkey E), pooled over bootstrap resamples. Tuning diagrams above illustrate four values of field shift (x-axis). (bottom) Analogous receptive field alignment for relational units (N=646 for Monkey P and 153 for Monkey E). Zero receptive field shift means the preferred relational orientation is parallel to the 1-object preferred direction vector. Each unit contributes twice by construction because relational orientation is undirected. **(f)** Firing rates of idealized retinotopic and relational units with zero receptive field shift. Each dot represents the 1-object receptive field location of an idealized unit. Hue represents firing rate in response to the stimulus (left). **(g)** Decoding response saccades from activity of recorded units. (top) Hue shows decoder output value and green cross its maximum for three 1-object trials from the training set. Gray circles are object positions. (bottom) Same plotting style for three 2-object evaluation trials. **(h)** Response decoder from (g) shows repulsive bias on 2-object conditions. Axes are the same as Figure 1d. Response is measured as the nearest local maximum of the decoder likelihood to the cued object. Colored points show response decoder mean with 95% confidence interval of the mean. Gray points show the same result for a surrogate dataset in which the 2-object neural activity is the average of the constituent 1-object responses.

We compared the models using each unit’s average firing rate during the stimulus and delay periods, before the cue appeared. For each unit, we used a subset of trials to estimate the preferred location, orientation or distance under the corresponding model. We then tested which model most reliably predicted responses of held-out trials (see Methods). In both monkeys, units tuned to the orientation connecting the two objects were most abundant (Fig. 3b). This result held in both brain areas, with relational orientation scored highest for 64–77% of units in each area and monkey (Supplementary Fig. 1), and for both stimulus and delay phases, with relational orientation scoring highest for 55-79% of units in each phase and monkey (Supplementary Fig. 2). In contrast, no unit was significantly better explained by the relational distance model (Fig. 3b). These results establish a strong tuning to inter-object orientation in both FEF and SEF during both stimulus presentation and working memory. Henceforth, we use “relational” to refer specifically to this relational orientation code.

#### Efficient coding

The prevalence of relational tuning raises a functional question: Why should a spatial population represent both individual object locations and the relations between them? Efficient-coding theory proposes that, under limited representational resources, neural populations should allocate their responses to maximize discrimination among behaviorally relevant stimuli^25–29^. Accordingly, we asked whether a population combining retinotopic and relational units would discriminate two-object configurations more effectively than a same-sized population composed only of retinotopic units.

We first tested whether retinotopic and relational units provided complementary information by training separate decoders to distinguish two-object configurations from their responses (Fig. 3c). When the line connecting the objects changed its orientation, decoding was more accurate from relational units than from retinotopic units. Conversely, when the objects occupied different locations but the connecting line retained the same orientation, decoding was more accurate from retinotopic units. A broader analysis of the two models showed that the changes in object configuration best discriminated by one were approximately orthogonal to those best discriminated by the other (Supplementary Fig. 3), suggesting a potential advantage to combining the two codes.

We next tested how the balance of retinotopic and relational tuning affected population coding (Fig. 3d). We sampled sub-populations of our neural data containing the same number of units but different proportions of relational and retinotopic tuning. We trained a decoder to recover the stimulus object locations from each sub-population’s activity. Decoding was most accurate for sub-populations with a balance similar to that of the full recorded population. Shifting this balance toward either retinotopic or relational tuning reduced decoding accuracy. Thus, the balance of relational to retinotopic tuning we observed in the brain was near optimal for representing the two-object configurations in our task.

#### Alignment of single-object and relational tuning

Many units were spatially modulated on both one- and two-object trials (692 in Monkey P, 183 in Monkey E, see Methods), allowing us to examine how the tuning for one-object stimuli relates to tuning for two-object stimuli. For retinotopic units, preferred locations were aligned across one and two-object stimuli (Fig. 3e, top). For relational units, the preferred orientation of the line connecting the two objects tended to align with the direction from fixation to the preferred single-object location (Fig. 3e, bottom). For example, a relational unit that preferred a single object above fixation tended to prefer two objects connected by a vertical line. Thus, although these units encoded different spatial variables across conditions, their single-object and relational preferences shared a common orientation.

#### Alignment of single-object and relational tuning predicts repulsion

The monkeys’ reports of the cued object location were biased away from the un-cued object, especially when the objects were close together (Fig. 1d). The alignment between single-object and relational tuning suggests a possible neural basis for this repulsion. Consider two objects positioned 90° apart on the ring (Fig. 3f, left). These objects strongly activate relational units tuned to the orientation of the line connecting them. Because of the observed alignment, these units’ single-object preferred directions lie parallel to that line, forming two opposite peaks displaced away from the actual object locations (Fig. 3f, right). If downstream circuits read out this activity according to each unit’s single-object preference, these signals would push the estimated locations apart. By contrast, overlapping retinotopic responses would pull the estimates toward one another (Fig. 3f, middle). The balance of these contributions could therefore determine whether remembered locations attract or repel.

To test this prediction, we trained a decoder to predict saccade direction on one-object trials using a pseudo-population randomly sampled from our recorded units but corrected to have unbiased receptive field locations (see Methods). On one-object trials, the decoder accurately predicts saccade direction (Supplementary Fig. 4). We then applied the decoder unchanged to two-object trials, where retinotopic and relational units yield attractive and repulsive biases, respectively. The combined activity produced two decoded peaks near the object locations, each displaced away from the other object (Fig. 3g). This repulsion was strongest when the objects were closest together (Fig. 3h), matching the dependence on object separation observed in behavior (Fig. 1d). Decoded repulsion was larger in Monkey E, consistent with its stronger alignment between single-object and relational tuning (Fig. 3e). In both monkeys, the decoded bias exceeded the behavioral bias. This difference suggests that downstream circuits may give relational units less weight than our decoder does.

### A continuum between retinotopic and relational tuning

So far we have treated retinotopic and relational tuning as two separate classes, but individual units showed intermediate response patterns. These patterns have a simple geometry when each preferred two-object configuration is drawn as a line connecting the objects (Fig. 4a). For retinotopic tuning, the lines converge near a common endpoint: one object occupies the unit’s preferred location while the other can occupy different locations. For relational tuning, the lines are approximately parallel: the objects can occupy different locations, provided the orientation connecting them is preserved. Intermediate units show partial convergence, with endpoints spread around a preferred location and lines more closely aligned in orientation.

**Figure 4.**
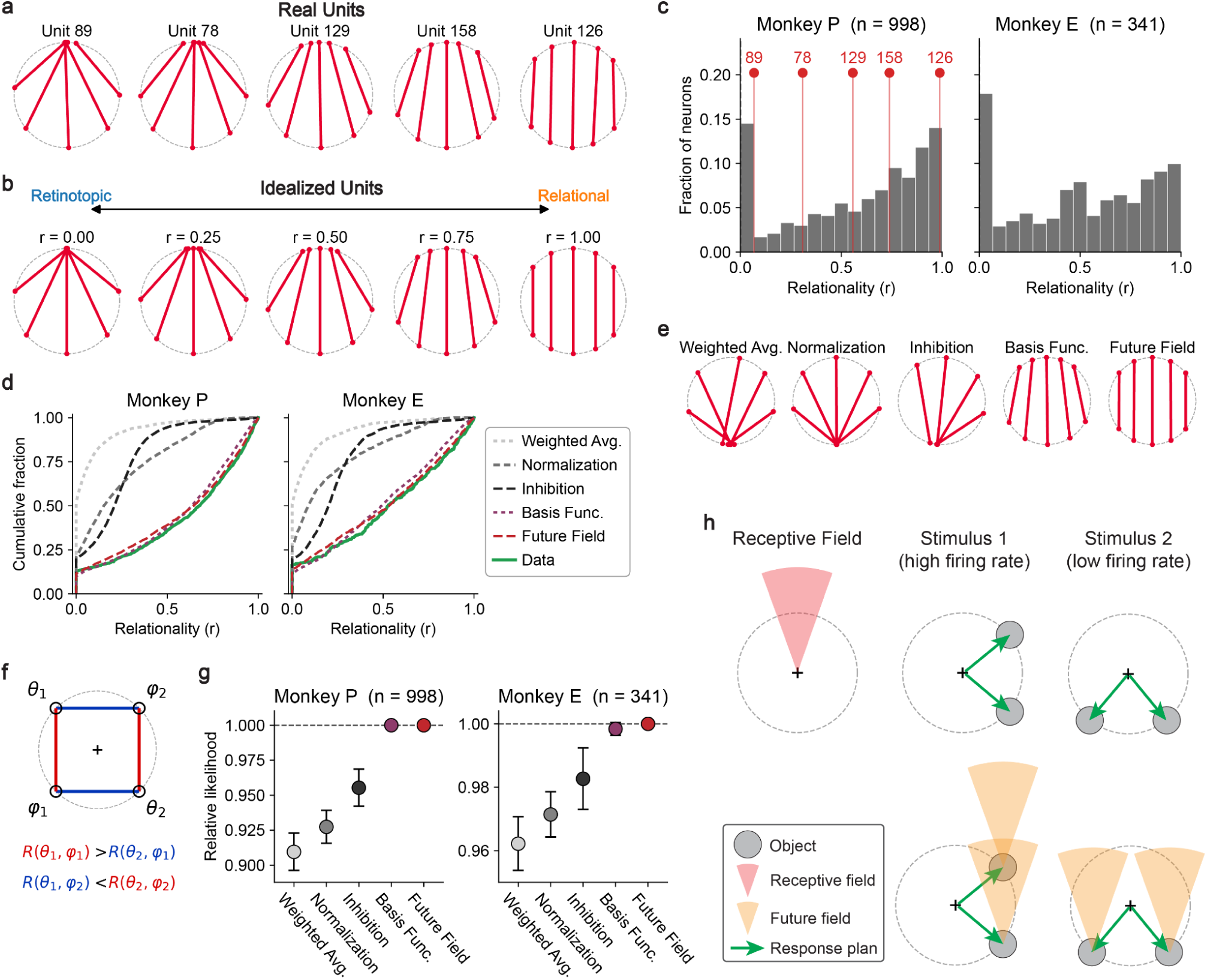
Relational spectrum and computational model. **(a)** Example units showing a relational spectrum, arranged left-to-right from retinotopic to relational. Unit 126 (far right) is also shown in Figure 2b. In each plot, chords show all stimuli eliciting locally maximal firing rates for 2-object stimuli with objects 80° apart, 130° apart, and 180° apart. Each plot is re-oriented to have comparable alignment. **(b)** Idealized plots corresponding to different values of the relationality metric (0 to 1 from left to right). Chords show the stimuli expected to elicit locally maximal firing rates among 2-object conditions with objects 80° apart, 130° apart, and 180° apart, given the definition of the relationality metric (see Methods). **(c)** Distribution of relationality across units. Red lines and labels show the values of the units in panel (a). **(d)** Cumulative distribution of relationality of the neural data (green) and synthetic data generated by Poisson samples of firing rates predicted by each fitted model (dashed lines). **(e)** Model fits to Unit 126 from panel (a). Plot style is the same as panel (a), except each column uses the firing rates generated from a different fitted model. **(f)** Schematic of an idealized relational unit’s responses to four 90-degree stimuli. Same style as Figure 2b. Equations show inequalities of response firing rates to these stimuli. **(g)** Model likelihoods. Each point is the mean over neurons of cross-validated per-trial Poisson likelihood of each model, normalized by that of the future field model. Errorbars show 95% confidence intervals of the mean. A constant null model has a relative likelihood of 0.64 for Monkey P and 0.84 for Monkey E. Dashed line shows equal likelihood as the future field model. **(h)** Future field model schematic. (top-left) Hypothetical angular receptive field (red). (middle column) Stimulus (top) with objects (gray dots) spanning a vector parallel to the receptive field direction. Green arrows show planned response saccades. Applying the response plans to the receptive field (bottom) yields two future receptive fields (orange), one of which contains an object. (right column) For a stimulus that is misaligned, neither future receptive field contains an object.

We developed a relationality index, r, to locate each unit along the spectrum from retinotopic to relational (Methods). Varying r from 0 to 1 generates tuning patterns ranging from purely retinotopic to purely relational, with intermediate values reproducing the geometry observed in hybrid units (Fig. 4b). We fitted r to each unit’s responses across the full stimulus space, and verified that it robustly captured the tuning properties of individual neurons (Supplementary Fig. 5). Relationality spanned the full range, with a distinct peak at 0 and a gradual increase in abundance across intermediate values toward a second peak at 1 (Fig. 4c), supporting a continuum between retinotopic and relational tuning.

### A computational model for relational tuning

What computational principles could account for the observed organization of spatial tuning? To address this question, we constructed models and tested which could account for the full range of tuning, from retinotopic through intermediate to relational. All models used a retinotopic receptive field, *E(.)*, described by a von Mises function—the circular analogue of a Gaussian—with preferred direction *μ* and tuning width *σ*. The models differed in how they combined or transformed responses to two objects at angles *θ* and *φ* to generate the two-object response, *R(θ, φ)*.

We first tested variants of three established multi-object models: contrast-weighted averaging^30–32^, divisive normalization^33–36^, and spatial summation of excitatory and inhibitory subfields^37–39^. In these formulations, each object contributes according to its retinotopic location. The averaging model weights and sums these contributions, the normalization model applies a saturating nonlinearity to their sum, and the inhibition model subtracts the contributions of an inhibitory receptive field:

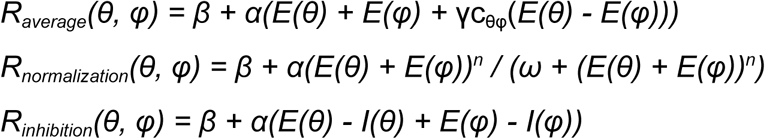

In all models, *β* is a baseline firing rate and *α* is the modulation amplitude. The averaging model weights the contributions of each object by a factor that depends on their relative contrast c_θφ_, scaled by a parameter γ controlling the strength of the contrast modulation (Methods). In the normalization model, *n* controls the nonlinearity and is shared across units, and *ω* controls strength. In the inhibition model, *E* and *I* describe excitatory and inhibitory spatial tuning. Each model had five parameters fitted per unit by maximizing the Poisson likelihood of the observed spike counts and was evaluated on held-out trials (Methods).

None of the three models adequately captured relational tuning (Fig. 4d,e). In these models, the conditional receptive field of one object given a fixed location of the other is constant. Placing the object at the mean of the receptive field maximizes firing rate wherever the other object is. In contrast, relational tuning requires a conditional preferred direction that depends on the conditioned object’s location (Fig. 2h). For example, consider a neuron tuned to vertical pairs (Fig. 4f). An object’s location is preferred because it lies directly above or below the other object. Moving the other object breaks that alignment, and the first object must move to restore it. The tested models cannot produce this dependence of spatial preference on the relationship between objects (Methods).

We next considered a basis-function model previously proposed to explain object-centered selectivity in SEF^40^. In this model, spatial responses depend jointly on retinotopic location and object orientation. We adapted it by multiplying the summed retinotopic responses by a factor tuned to the orientation of the line connecting the two objects:

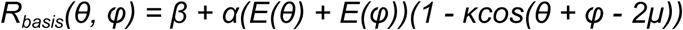

Here, *κ* controls the strength of orientation modulation. An object at the preferred retinotopic location can now evoke a weaker response than an object elsewhere if the latter forms a better-aligned pair with the second object. This interaction allows the preferred location of one object to change with the location of the other. With five fitted parameters per unit, matching the preceding models, the basis-function model better captured the observed distribution of relationality (Fig. 4d,e) and achieved higher cross-validated likelihood across the population (Fig. 4g). Thus, modulating retinotopic responses by inter-object orientation allowed the model to capture the range of tuning from retinotopic to relational.

A key observation in the oculomotor system is that neurons can respond to stimuli that will enter their receptive fields only after an impending eye movement^11,12^. This motivated us to develop a “future-field” model that extends this anticipatory response to conditions with two objects and two potential saccades (Fig. 4h). In the model, a neuron’s response is influenced by both its receptive-field response at the current fixation, *E_present_*, and its responses predicted after saccades to each object, *E_future_*. After a saccade to one object, the retinal position of the other is its displacement from that object. The future component therefore provides relational tuning through the anticipated change in gaze. We model each unit as a weighted average of *E_present_* and *E_future_*.

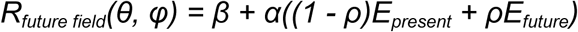

Here, *ρ* controls the relative contribution of predicted future responses, ranging from purely present-field responses at *ρ = 0* to purely future-field responses at *ρ = 1*. The future-field model also had five fitted parameters and performed similarly to the basis-function model both in terms of likelihood and the cumulative distribution of relationality (Fig. 4d,e,g). These results held in both FEF and SEF (Supplementary Fig. 1). By anticipating future saccades, this model makes the preferred location of one object depend on the location of the other. This serves the same role as orientation-dependent modulation in the basis-function model.

Together, these results show that the observed continuum can arise from a common computational framework in which spatial preferences depend on the configuration of the pair. Both orientation-dependent modulation and prediction of future gaze positions can generate this dependence, accounting for forms of tuning that the averaging, normalization, and inhibition models could not capture.

## DISCUSSION

How does the brain represent scenes with multiple objects? Does it represent objects independently or relative to one another^41,42^? Our findings show that, during a spatial working memory task, some neurons in FEF and SEF showed retinotopic responses that treated objects independently but the majority of neurons instead encoded relative object locations. The retinotopic and relational codes provided complementary information for distinguishing two-object stimuli. Furthermore, relational neurons had 1-object receptive fields that aligned with the relative positions of their preferred two-object stimuli, and this alignment predicted systematic biases in remembered locations.

The prevalence of relational tuning in individual FEF and SEF neurons is striking given decades of work characterizing these areas through receptive fields defined by stimulus location relative to gaze direction or head position. Why did earlier studies not reveal such prevalent relational tuning? We suspect that both task design and the prevailing conceptual frameworks contributed. Tasks emphasizing a single spatial location offered limited opportunities to reveal relational tuning, while the emphasis on eye- or head-centered position codes encouraged interpreting contextual effects as modulations of this code.

There were, however, indications that spatial selectivity could depend on relationships between objects. Studies in SEF identified neurons selective for the left or right side of a reference object across changes in retinal position and visual form^17–19,43^. More recent work showed that landmarks bias both the receptive fields of neurons in FEF and SEF and the reference frame in which other targets are encoded, indicating that relative landmark-target position modulates responses^14–16,44,45^. Our findings extend these observations from categorical object-relative positions and landmark effects to tuning for inter-object orientation across continuously varying configurations. Moreover, this tuning emerged in a task requiring memory for individual locations of multiple objects without an explicit judgment of their relationship or an asymmetry in the role of the objects. We found that relational tuning was widespread in both FEF and SEF. These findings invite a broader view of these areas: Their spatial representations can organize multiple locations in relation to one another, extending their established roles in selecting and maintaining individual saccade targets. This organization may help link the spatial structure of a scene to the actions available within it.

Efficient coding provides a broad principle for understanding how neural populations use limited resources to distinguish relevant stimuli^25–27^. Our findings extend this principle to spatial coding in FEF and SEF: Retinotopic and relational units contributed complementary information, and their observed balance supported near-optimal decoding of the two-object stimuli in our task. This organization may partly reflect extensive training on our task. The natural demands on FEF and SEF are considerably richer. During active vision, we make multiple saccades per second, repeatedly selecting objects within scenes containing many potential targets. Each movement changes objects’ retinal locations while preserving their arrangement relative to one another. Representing both locations and relationships could therefore be particularly useful for circuits that guide gaze through such scenes. Whether and how our efficient coding result extends to settings with more objects and freely chosen gaze shifts remains to be explored.

We found that relationally tuned neurons had receptive fields in single-object conditions that were aligned with their preferred inter-object orientation in two-object conditions. This alignment provided a neuronal account of a counterintuitive, longstanding psychophysical effect^21,22^, which we also found in our monkeys’ behavior: Object positions are repelled away from each other in working memory of two-object scenes. A readout trained to localize single objects from neural activity in our data produced this repulsion when applied to two-object responses, including the stronger bias at smaller separations observed in the monkeys’ behavior. While this result suggests that relational neurons impact motor execution, whether the contribution to motor execution differs between retinotopic and relational neurons remains unknown. The larger repulsion in our decoder than in behavior suggests retinotopic neurons may contribute more to motor execution than relational neurons. Future experiments could identify the projections and effective weights through which the two codes guide behavior and explore whether those weights change with task demands.

We found a continuum from retinotopic to relational tuning across recorded units, which provides a strong constraint on the computations underlying multi-object coding. The averaging, normalization and tuned-inhibition models we tested could not explain relational tuning, which the basis-function and future-field models captured. The basis-function account requires aligned retinotopic and orientation-selective inputs to interact multiplicatively, a circuit arrangement that remains to be established. The future-field account links relational tuning to a well-established phenomenon in the oculomotor system: Shortly before a saccade, neurons can represent where stimuli will fall on the retina after the eyes move^11^, supported by corollary-discharge signals conveying information about the impending movement^13,46^. Receptive-field shifts have also been observed during delayed saccade preparation, extending spatial reorganization beyond the immediate perisaccadic period^47^. Our model suggests that anticipating multiple potential gaze shifts could contribute persistently to relational tuning. Recent studies have debated whether remapped receptive fields undergo a convergence effect that biases them towards the saccade endpoint^48,49^. Our results are agnostic to this debate, and the future field model produces relational responses to two-object stimuli both with and without convergent remapping (Supplementary Fig. 6).

These models leave open where relational signals arise and how they are implemented. The predictive responses proposed by the future-field model could draw on corollary-discharge pathways, local recurrent circuitry or interactions between them^13,46,47,50^. Relational signals could also be inherited from elsewhere. Candidates include the parietal area 7a, where object-centered responses have been reported^51^, and the medial temporal lobe, where relational representations of environmental structure have been studied^52–57^. Recording and perturbing interconnected areas during a multi-object working memory task could reveal whether relational tuning is constructed within frontal oculomotor circuits, inherited from other regions, or generated through interactions across them.

Our task involved two potential targets within a constrained spatial arrangement: Both objects lay on a ring at a fixed eccentricity. Allowing their eccentricities to vary would reveal whether the same tuning principles extend to a broader range of object positions and separations. With three or more objects, the representational problem also changes: Multiple relationships coexist, raising the question of which relationships individual neurons encode. Some units might be selective for particular pairs, whereas others might combine several relationships or respond to larger configurations. The future-field framework could extend by integrating over additional potential saccades, while grouping and attention could determine which objects contribute to each unit’s response. Exploring these richer settings may reveal a broader repertoire of representations for multi-object scenes.

Our task also leaves unresolved the contributions of covert attention, working memory, and motor planning to relational tuning^58–60^. Because either object could become a saccade target, maintaining both locations may involve preparing two potential eye movements. The future-field model suggests how these functions could interact: Anticipating a gaze shift to one remembered object represents the other relative to it. Whether relational tuning depends on such prospective movements could be tested by varying the required response while keeping the memory demands unchanged, or by revealing the response modality only after the delay. These experiments would help establish whether the relational code reflects the arrangement held in memory, the actions available within it, or an interaction between the two.

Our analyses focused on spatial tuning, but successful performance on our task also required retaining the association between each object’s identity and its location^61^. Object features are represented widely in cortical circuits including the inferotemporal, parietal, and ventrolateral prefrontal cortex^1,62–64^, and individual prefrontal neurons can jointly encode object identity and location during working memory^65^. Relational coding may facilitate this binding: A relational spatial representation in FEF and SEF may allow a binding representation elsewhere to have less spatial precision or lower-capacity inter-area communication. In a network model of our task, we found that a capacity-constrained binding pool communicates preferentially with relational units over retinotopic units (Supplementary Fig. 7). This result motivates testing whether inputs carrying object identity preferentially engage relational units in FEF and SEF, and whether these interactions help preserve object–location associations as memory demands increase.

The discovery of abundant relational receptive fields in frontal oculomotor cortex connects a broad cognitive question “how does the brain represent a scene?” to the tuning of individual neurons. It brings scene representation into a common framework with efficient coding, behavioral biases and computations that anticipate future gaze shifts. These connections open a research program spanning cells, circuits, and cognition: how relational tuning emerges from interactions among neurons, how it organizes representations of scenes, and how those representations guide memory and action. Relational receptive fields provide a concrete foothold for understanding how the brain builds a structured representation of the world.

## MATERIALS AND METHODS

### Experimental procedures

All experimental procedures conformed to the guidelines of the National Institutes of Health and were approved by the Committee on Animal Care at the Massachusetts Institute of Technology. We performed all behavioral and neural experiments on two rhesus monkeys (Macaca mulatta). Monkey P was female, 10 years old, and weighed 8kg. Monkey E was male, 10 years old, and weighed 12kg. During all experiments the monkeys were seated comfortably in a primate chair and head-fixed in a dark room. The task was displayed on an Acer H236HL LCD monitor (23 inches diagonal size, 60 Hz refresh rate, 1920 by 1080 pixel resolution) positioned 22 inches away from the monkey’s eyes. The pupil position of the right eye was recorded at 1kHz by an infrared camera (Eyelink 1000, SR Research Ltd., Ontario, Canada). A mapping from pupil position in Eyelink camera coordinates to gaze point on the screen was calibrated at the beginning of every session. To do this calibration, the monkey was trained to fixate sequentially on each point in a 3×3 grid spanning the height and width of the display. The intervals of fixation were extracted and the eye positions during these intervals were used to fit a linear regression from pupil position to screen coordinate.

### Behavioral Task

The task was implemented in MOOG^66^ and MWorks (http://mworks-project.org).

#### Fixation phase

A square arena with sidelength 24° appeared with a grayscale pink noise background with 1/f power spectrum, normalized to 25% maximum screen brightness. The background was randomly sampled each trial but unchanged within a trial and served to minimize afterimage effects that could be used for visual working memory. In the center of the arena was a white fixation cross with diameter 0.94°. If the gaze position was within 3.36° of the center of the fixation cross, the monkey was considered to be fixating. One of our monkeys had difficulty maintaining fixation, so we introduced a variable fixation cross size for both monkeys at all times the monkey was fixating: If the gaze position was further than 1.44° from the center of the fixation cross, the fixation cross grew in size linearly proportional to the gaze eccentricity passed through a 100ms boxcar filter. After holding fixation for 400ms, the task progressed to the stimulus phase.

#### Stimulus phase

Objects appeared on a ring of radius 8.4° around the fixation point. Each object was a white circle of radius 2.06° with an image of a fruit occupying most of the circle. During physiology sessions, the fruit was randomly sampled without replacement from a set of three images {apple, blueberry, orange}. The fixation cross remained through this phase. After 1000ms, the task progressed to the delay phase unless the monkey broke fixation.

#### Delay phase

The objects disappeared. After 1000ms, the task progressed to the cue phase unless the monkey broke fixation.

#### Cue phase

The cue was a white square of sidelength: 4.12° presented at the center of the screen, occluding the fixation cross. The square contained one fruit randomly selected from those in the current trial’s stimulus. After 180ms, the monkey was allowed to make a saccadic response, provided fixation was maintained until that point.

#### Response phase

250ms after cue onset, a visual change occurred that helped indicate the monkey to respond. The cue became partially transparent (alpha 0.59) and a foveal window appeared. The foveal window was a circle with radius 2.4° that moved with the eye such that the center of this circle was the monkey’s gaze point. The foveal window effectively created a “hole” in the background noise, through which the monkey could see the objects hidden beneath. This was a useful tool during training, because it allowed the monkey to search for the correct stimulus object without using peripheral vision. The foveal window had a smooth boundary, with full background transparency within 1.2° and a linear radial gradient of opacity to full background opacity at 2.4°. The foveal window did not allow the monkey to view any stimulus objects after the cue onset before making a response saccade, hence did not allow the monkey to cheat on the task. The foveal window also did not give rise to any unusual eye movement behaviors. The monkeys made saccadic responses with normal eye velocity and eccentricity. The endpoint of a response saccade was detected if the gaze position was no longer within 3.36° of the center of the screen and the gaze velocity was below 7.2 degrees per second. In practice, these parameters detected saccadic responses very reliably for both monkeys.

#### Feedback phase

After the monkey responded, all objects were revealed with partial opacity (alpha 0.59) for visual feedback. This feedback lasted 500ms for correct trials and 1000ms for incorrect trials. During the feedback phase the monkey received a water or juice reward for correct trials. A response was deemed correct if it lay within 2.4° of the center of the target object. The reward was delivered via a tube next to the monkey’s mouth, and the flow of reward was controlled by a solenoid. The reward scaled with trial difficulty: The monkey received 75% more reward for correct 2-object trials than correct 1-object trials. The reward also scaled with accuracy, with maximum reward for a response position exactly the same as the center of the target object. Reward scaled linearly down to zero if the response was more than 2.4° from the center of the target object. Upon a correct response, in addition to reward, the monkey received an auditory cue (high-pitched ding). Upon an incorrect response, the monkey received a different auditory cue (low-pitched honk).

#### Inter-trial interval

A gray screen (luminance 0.2 times the maximum brightness of the screen) was presented for 500ms after correct trials and 1000ms after incorrect trials before the fixation phase began for the next trial.

If the monkey’s gaze position was greater than 3.36° from the center of the screen at any point until the response phase, the monkey was considered to have broken fixation. If the monkey broke fixation during the fixation phase of the task, the trial did not restart and the task waited for the monkey to regain fixation. If the monkey broke fixation during the stimulus, delay, or cue phase, the trial aborted. In this case, as soon as fixation was broken, the arena turned into a maroon square (RGB value [102, 51, 51]) for 1000ms, followed by a 1000ms ITI, followed by the onset of the next trial. Simultaneously, the monkey got auditory feedback (same low-pitched honk as for incorrect trials).

We randomly interleaved 1-object trials with 2-object trials. For each trial, with probability 0.75 we delivered a 2-object stimulus. The two objects’ positions were randomly sampled on the ring subject to the constraint that they were no less than 10.8° visual angle apart (80° arc-angle around the ring). With probability 0.2 we delivered a random 1-object stimulus in which one object was randomly uniformly sampled with eccentricity 8.4°. With probability 0.05 we delivered a left/right 2-object stimulus. These left/right trials were excluded from further analysis to minimize spatial biases in results.

All analyses used completed trials only, namely trials for which the monkey did not break fixation and hence generated a response.

### Neurophysiology

Each monkey underwent a first surgery under general anesthesia to attach three titanium headpins to the skull for head fixation and a second surgery to implant a medial chamber of polyether ether ketone for neurophysiology recording. A craniotomy was made over SEF in each monkey. We recorded simultaneously from right-hemisphere dorsomedial frontal cortex (SEF) and left-hemisphere frontal eye field in both monkeys. We recorded from a 4mm x 6mm patch of superficial SEF in each monkey, randomly sampling recording sites within this patch during each session. In the frontal eye field, we recorded from the lateral bank of the superior arcuate sulcus and the anterior bank of the genu of the arcuate sulcus, randomly sampling recording sites from a 5mm x 8mm patch of cortex during each session (Supplementary Figure 8a).

We recorded from SEF using non-reinforced 10mm Neuropixels 1.0 probes^24^. Because these probes are very short and fragile, we developed a custom guide tube system (Supplementary Figure 8b). This guide tube system allowed us to penetrate the primate dura with a very short (2mm) tip of the guide tube and insert the Neuropixels probe through the guide tube into SEF via a micromanipulator, while controlling the stereotaxic coordinates of the insertion.

We recorded from FEF using Plexon V-probes with 64 channels and 50 micron inter-contact spacing. We used V-probes to record from FEF because both monkeys had pre-existing chambers centered over the frontal midline for SEF and did not lie over FEF, so we had to approach FEF through an angled trajectory that was too long for 10mm Neuropixels probes. We used a custom angled grid and guide tube system for FEF recordings that was compatible with simultaneous SEF recording.

We recorded Neuropixels data at 30kHz frequency with SpikeGLX software and V-probe data at 30kHz frequency with OpenEphys software. We recorded 23 total sessions (11 from Monkey P and 12 from Monkey E).

### Data Processing

Neural data were temporally synchronized to behavioral data by a linear regression based on a sync pulse sent at the beginning of every trial, and the timing of task events on the monitor was adjusted by a display delay measured via a photodiode in the corner of the monitor.

Neural data were then spike sorted using Kilosort 2.5. For V-probe data, spike sorting results were curated manually with a spike sorting curation gui (Phy), through which we did splitting, merging, and labeling of the units. For Neuropixels data, we performed motion estimation and correction using MEDiCINe and SpikeInterface ^67,68^. The quantity of units from the Neuropixels data was prohibitively large for manual merging and splitting, so we used a custom automated pipeline to do splitting based on unit statistics, then manually labeled each unit to identify putative single-units and multi-units. Our pipeline also detected and excluded duplicated units, which have the potential to give spurious modeling results. Broadly, we were extremely conservative with our spike sorting: We thoroughly examined a variety of statistics and plots for every unit, only labeled units as single neurons if they were very clearly single neurons, and liberally classified as noise anything that did not have a clear neuron-like waveform. We included units labeled as multi-units in all of our analyses. We found no evidence that our results were qualitatively impacted by the inclusion of multi-units: Restricting to single neurons showed qualitatively similar results (Supplementary Fig. 9).

After spike sorting, we identified for each recorded unit an interval of time for which the unit was stable by running a filter on firing rate, thresholding the derivative of firing rate over time to identify points where the unit was lost or gained, and validating manually. For many units the identified interval of stable recording was the entire session, but some units were lost or gained partway through the session. We excluded all spikes outside of this stable range.

We also excluded from analysis all units that were not stable for at least 300 completed 2-object trials, and all units with mean delay-phase firing rate less than 1Hz. We also excluded all units that had no significant modulation across 2-object trials, as defined by the following model-agnostic criterion: Under 5-fold cross-validation with 10 bootstraps, we measured the Poisson deviance explained by a Gaussian kernel smoother with standard deviation 30° over objects. We excluded all units for which this deviance explained was less than 0.02 or was not statistically significant. This excluded 1,672 of 3,011 recorded units. Upon visual inspection, these excluded units had no discernable modulation with respect to the stimulus.

### Retinotopic, Relational, and Proximity Models

Each model has a single parameter. For the retinotopic model this is the preferred direction. For the relational model this is the preferred orientation. For the proximity model this is the preferred inter-object distance. We can formalize this in equations. Suppose an anchor object has angle θ. Then the preferred direction(s) μ(θ) are given by:

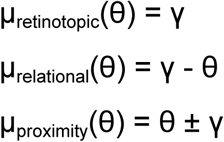

where γ denotes each model’s parameter, fitted to each unit. We fit γ for each model alone, without creating or fitting any other parameters. In particular, we made no inference or assumptions about receptive field size or shape, baseline rate, or modulation amplitude. The intuition is that each model predicts an ordering on responses that share one object position. Consider trial A with objects at (θ, φ_A_) and trial B with objects at (θ, φ_B_). These trials share an object at angle θ. Each model (given a value of its parameter γ) then predicts whether trial A or trial B elicits a higher firing rate:

- Retinotopic model: Trial A will fire more if and only if φ_A_ is closer to the preferred direction than φ_B_.
- Relational model: Trial A will fire more if and only if the angle of the vector between the objects in trial A is closer to the preferred orientation than the angle of the vector between the objects in trial B.
- Proximity model: Trial A will fire more if and only if the distance between θ and φ_A_ is nearer to the preferred inter-object distance than the distance between θ and φ_B_.

Only these relative firing rate predictions are used for model-fitting.

We needed a way to handle spatial smoothness, because object locations were sampled continuously around the ring. Consider a pair of two-object trials with anchors at locations θ_A_ and θ_B_, we defined the pair’s weight to be:

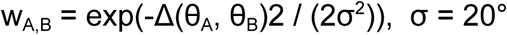

where Δ is angular distance. We use these weights as coefficients on the model’s loss function, so the loss is driven by trials that have nearby anchors. We optimize each model to maximize its accuracy

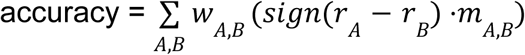

where m_A,B_ is the model’s signed ordering of the firing rates for trials A and B (assuming an anchor at the midpoint of the two trials’ anchors) and r_A_ is the neural firing rate on trial A. Because we only recorded a finite number of trials, we used search to compute the value of the model’s parameter that maximized accuracy.

We used five-fold cross validation, evaluating the model on the held-out fold for the results in Figure 3b. We computed the win probability of each model by 1000-fold bootstrapping of held-out trials. A unit contributed to the barplot only if its held-out accuracy was higher than that of both other models for at least 950 of 1000 bootstraps.

### Efficient Coding

For each neuron, let z_retinotopic_ and z_relational_ denote the held-out accuracy of the retinotopic and relational models on that neuron, respectively, from Figure 3b. For Figure 3c, we selected the top 10 neurons by z_retinotopic_ from each monkey and used the resulting 20 neurons as the retinotopic population (blue), and computed the relational population (orange) analogously. These neurons came from different sessions, so for each neuron and each stimulus, we computed the neuron’s response for all trials where both objects were within 15° of the stimulus object position. We then used 5-fold cross-validation on these trials, fit a logistic linear decoder to discriminate the two stimuli from single-trial population vectors sampled from the train trials, and evaluated it on single-trial population vectors sampled from the test trials.

The insets in Figure 3d show the space of z_retinotopic_ (x-axis) and z_relational_ (y-axis). We ranked each neuron by z_relational_/z_retinotopic_ and sampled sub-populations without replacement from an exponential tilt distribution exp(λ rank) on this rank. We swept λ so that the mean ranks were evenly spaced from ∼0.1 to ∼0.9. The x-axis values in Figure 3d are the empirical mean ranks. We also ensured that each sub-population had the same number of neurons (which was the maximum number possible for each monkey given the minimum and maximum λ). We used 100 sub-population samples for each λ.

To decode the stimulus, we used a feedforward MLP with two hidden layers of 64 units and a 96-unit output layer with ReLU activations. We trained it by gradient descent with the Adam optimizer with batch size 128, and learning rate 2×10^-3^ for 1,000 steps. The target was a sum of two Gaussian bumps (standard deviation 12°) centered at the object positions on a 96-unit ring of units.

The input to this decoder was a pseudo-population of units with de-biased preferred direction distribution. For each unit in the sub-population, we sampled a random angle to re-orient all of its responses, thereby eliminating any correlation of preferred directions the population may have had, which may have confounded our results. Also, since the units came from different sessions, to compute their response for a given stimulus we took an average of the response over recorded trials, weighted by the similarity to the given stimulus (Gaussian with standard deviation 10° on each object), then applied Poisson sampling. To evaluate decoder error, we computed the two highest local maxima of the decoder’s output, and computed the mean angular difference between each stimulus object and the nearest such local maximum.

### Response decoding

For a unit to contribute to Figure 3e-h, it needed to have a measurable 1-object receptive field orientation. To evaluate this, we excluded units that were not recorded for at least 100 1-object trials. We then fit a 1-object preferred direction to each unit given responses on 1-object trials. Specifically, this preferred direction was optimized to maximize the correctness of the resulting ordering of firing rates on 1-object trials, analogous to our fitting procedure for the 2-object models in Figure 3b. If this fit was not statistically significant (p < 0.05 from a shuffled null), we excluded the unit. We estimated the 1-object receptive field direction from 50 bootstrap resamples with replacement of 1-object trials, and allowed each bootstrap to contribute to Fig. 3e. We also needed to estimate the preferred direction/orientation for 2-object retinotopic/relational models. We did this in an analogous way: For each plot, we rejected units whose firing rate rank was not significantly different from a shuffled null, and used 50 bootstrap resamples to estimate the preferred direction/orientation. Note that for the retinotopic histograms, each bootstrap contributes one point, whereas for the relational histograms, each bootstrap contributes two points (exactly 180 degrees apart) because we do not prescribe a directionality of the relative vector between the objects. We further tested whether these histograms were significantly different from a uniform distribution using a Kuiper statistic with one value per neuron, no duplicated points, and 10,000 Monte Carlo draws. We found that all were significantly different from a uniform distribution (p = 0.0001 / 0.0003 for Monkey P retinotopic/relational and p = 0.0001 / 0.0004 for Monkey E retinotopic/relational).

In the schematic in Figure 3f, 300 idealized retinotopic and 300 idealized relational units are shown. Each has zero receptive field shift between 1-object and 2-object preferred directions. Each has a von Mises receptive field with half-width 60°. Each is plotted at its 1-object receptive field location.

Figure 3g,h use recorded units. For each monkey, we performed 5 bootstraps. For each, we sample 100 units with replacement and without bias. Receptive field locations of our recorded units are not uniformly distributed, and that non-uniformity may impact our readout, so to combat this we used 128 clones of each unit, where each clone was rotated and reflected in a different orientation to evenly tile the ring. These clones used the real responses of the unit (no simulated data was used), just rotated and reflected the stimulus systematically. This gave us a population of 12,800 units to fit a decoder with no bias in receptive field location.

We then fit a decoder on this population response. The decoder is a linear readout onto a ring of output channels, and each unit contributes a truncated cosine lobe centered on its 1-object preferred direction. For each unit u we fit two parameters: A preferred direction μ_u_ and an amplitude a_u_. Then the decoder value at angle θ is

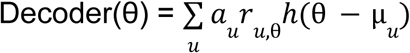

where h() is a positive rectified cosine and *r_u_*_,θ_ is the response of unit u to a 1-object stimulus at angle θ (computed as a mean of responses to all trials weighted by a circular-Gaussian kernel with scale 15° centered at θ). The amplitudes and preferred directions were fit jointly by alternately solving for amplitudes and sweeping preferred directions. Preferred directions were initialized using a cosine fit.

The decoder was then applied unchanged to 2-object conditions, given the same bootstrapped sample of neurons. To compute the response error for an object, we measured the decoder response to each object as the nearest local maximum of the decoder output to the object position.

For the null decoding (gray points in Figure 3h), we applied exactly the same methodology, except that neural response for each 2-object condition (θ, φ) was the average of neural responses for the two 1-object conditions with objects at θ and φ.

### Computational Models

Each model had 5 parameters: Baseline β > 0, amplitude α > 0, preferred direction μ, tuning width σ > 0, and one model-specific parameter. Each model had an excitatory receptive field E defined by a von Mises function with mean μ and width σ

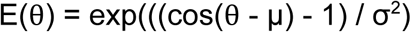

We now detail the models.

#### Weighted Average

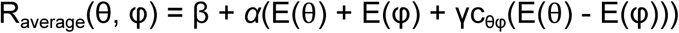

where

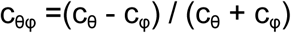

where c denotes the average luminance of an object (fruit image on white disc) relative to the background luminance. Here γ is the model’s fifth parameter, which controls how much luminance modulation affects the average and is constrained such that −1 < γc_θφ_ < 1.

#### Normalization

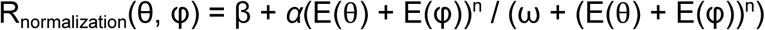

where ω > 0 is the model’s fifth parameter and controls the strength of the normalization. In the literature, typically n is chosen to be 1 or 2 ^34,35^. For the results in Figure 4 we let n = 1, though we explored n = 2,3,4,5 and found no significantly better performance for any of those values. Note also that the normalization model often considers the activity of a local population of neurons in the denominator, assuming some similarity in tuning across nearby neurons. We did not explore this direction and made the simplifying assumption that normalization happens at the single-neuron level (or within a population of identically-tuned neurons).

#### Inhibition

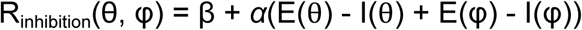

where I is an inhibitory receptive field defined by a von Mises function with mean ψ and width σ. The inhibitory receptive field mean ψ is the fifth parameter for this model. We share the width σ between the inhibitory and excitatory receptive fields. We also tested a version where the means of the inhibitory and excitatory receptive fields are the same and their widths are allowed to differ, but that performed worse. A large variety of models have been proposed in the literature that integrate excitatory and inhibitory fields ^37–39,69^. We chose this formulation because it is simple and matches the computational complexity of the other models, and we cannot exclude the possibility that a more flexible version of the inhibitory model (or of any of the other models) may fit our data better.

#### Basis Function

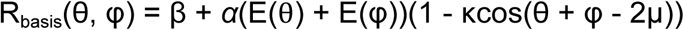

where κ > 0 is the fifth parameter and controls the strength of the multiplicative gain. Note that this formulation assumes the relational tuning (preferred angle of the relative vector between the objects) is aligned with the 1-object receptive field. This assumption was supported by our analysis of receptive field orientations (Fig. 3e) and allows the model to have the same complexity as the others. However, we cannot exclude the possibility that a version that allows independent 1-object and relational receptive field orientations may fit our data better.

#### Future Field

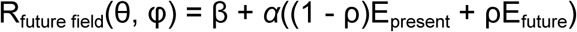

where 0 < ρ < 1 is the fifth parameter. Note that ρ = 0 corresponds to a retinotopic average response and ρ = 1 corresponds to a purely relational response (sensitive only to relative, not absolute, object position).

We fit each model independently to each neuron by maximizing the Poisson log-likelihood of the neuron’s responses with gradient ascent. Some models may produce negative rates under some parameters, in which case those rates are rectified to zero. For model comparison, for each neuron we used 10 random 80%/20% train/test splits (with splits shared across models) and measured Poisson likelihood on the test set.

### Relationality Metric

In panels b and c of Figure 4, we report results of a relationality metric in [0, 1]. We computed this metric on neural data and on synthetic data generated by Poisson samples of the fitted model-predicted firing rates. To compute the relationality metric, we fit the following model:

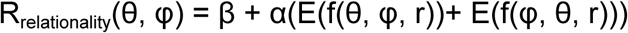

where f(θ, φ, r) is the angle of object θ with respect to the point (1 - r) * fixation + r * midpoint, where midpoint is the midpoint (in cartesian coordinates) of the two objects. In other words, the objects are integrated over a receptive field with respect to some origin that lies between fixation and object midpoint, and the relationality r in [0, 1] prescribes how close the origin is to the object midpoint. If r is zero, the origin is fixation and the response is purely retinotopic. If r is one, the origin is the object midpoint and the response is purely relational. To measure r per neuron, we use the same fitting procedure as for the models above.

## MATERIALS AND METHODS

### Figure 1

#### Panel c

Responses with a radius below 6° visual angle were deemed uncommitted and excluded from the plots. For Monkey P, 0.3% of 1-object and 1.6% of 2-object trials were uncommitted. For Monkey E, 2.7% of 1-object and 2.6% of 2-object trials were uncommitted.

#### Panel d

For each completed trial we recorded the angular positions of the objects, the index of the cued object, and the polar coordinates of the response saccade. We computed the response error as the angular difference between the response and the target (cued object position). Completed 2-object trials were grouped into nine 25°-wide bins of inter-object angle, centred at 80° through 280°. Trials with an absolute response error greater than 30° were excluded, since those may reflect lapses or responses to the non-target object.

#### Panel e

To distinguish spatial errors from swap errors, we modeled each session’s 2-object responses as a mixture of three components:

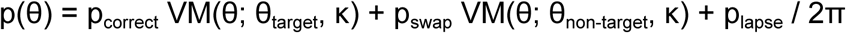

where VM is a von Mises density. Three parameters (p_correct_, p_swap_, κ) were fitted with expectation-maximisation, and p_lapse_ = 1 - (p_correct_ + p_swap_). Reported swap error is p_swap_. Reported spatial error is the circular standard deviation of the von Mises component given the fitted κ.

### Figure 2

#### Panels a-d

Each of the example stimuli in the left and right columns are prototypes for averaging. The firing rates for these are averaged over all trials where the objects were both within 15° angle (with respect to fixation) of these prototypes. The empirical mean thus deviated slightly from the prototypes, but this deviation was minimal (Supplementary Fig. 10). Like the behavioral analyses, all neural analyses considered only trials where the monkey did not break fixation, and hence gave a response. However, we did not condition or filter based on the accuracy or correctness of that response.

#### Panels e,g

Six anchor positions were selected evenly around the ring (5°, 65°, 125°, 185°, 245°, 305°). A 2-object trial contributed to an anchor if one of its objects was within 30° of the anchor. The angle of the other object is the non-anchor angle. Objects were un-ordered, so each trial contributes to two anchors (one for each object). For each anchor, firing rate was modeled as

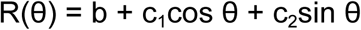

and b, c_1_, and c_2_ were fit with ordinary least squares. The preferred direction arctan(c_2_, c_1_) was deemed the receptive field center for that anchor.

#### Panels f,h

Note that a perfectly retinotopic unit corresponds to a horizontal line in the lineplot (right), and the preferred direction is the height of the line. A perfectly relational unit corresponds to a diagonal line with slope −1 (which wraps around because the x- and y-axes are angles), and the preferred relational orientation is determined by the intercept of that line.

### Figure 4

#### Proof that average, normalization, and inhibition models cannot capture relational tuning

The failure of the weighted average, normalization, and inhibition models to capture relational tuning can be formalized mathematically: All three models are of the form R(θ, φ) = f(g(θ), g(φ)) for some functions f and g, with f monotonically increasing in both arguments. To see that no model of this form can produce a relational response, note that for any pair of locations θ_1_ and θ_2_, the sign of R(θ_1_, φ) - R(θ_2_, φ) cannot depend on φ due to monotonicity of f. This implies that the conditional preferred direction must be constant, unlike relational units (Fig. 2g,h). This contradiction is most clearly illustrated by a relational unit tuned to vertically aligned stimuli and the four points {θ_1_, θ_2_, φ_1_, φ_2_} and 2-object stimuli shown in Figure 4f. In this setting, R(θ_1_, φ_1_) > R(θ_2_, φ_1_) whereas R(θ_1_, φ_2_) < R(θ_2_, φ_2_). Relational tuning thus requires more than a nonlinearity: It requires sign-reversal of the conditional response function, which is not possible by integrating receptive fields in a gaze-centered coordinate system with monotonic nonlinearities.

#### Panel a

For each inter-object angle (80°, 130°, 180°), we considered 180 stimuli with that inter-object angle (spaced at increments of 2° around the ring). We computed the firing rate for each of these sample trials by averaging the firing rate for all sufficiently similar trials, namely trials where the objects are within 15° of the sample trial. We then computed all local maxima of these firing rates and for which the difference in firing rate with the neighboring troughs was at least 10% the dynamic range of the neuron (to exclude local maxima arising from noise/jitter). For each local maximum, we plotted the segment between the objects.

#### Panel g

The y axis is mean cross-validated model likelihood over neurons. To compute this, for each neuron we performed 10 bootstraps with samples of 80% train / 20% test trials, fit the model on the train trials and evaluated Poisson likelihood on the test trials.

### Supplementary Figure 3

Figure 3c compares two perturbations of a particular stimulus and finds that retinotopic and relational sub-populations provide different directions of sensitivity. Supplementary Figure 3 shows this complementarity over the entire space of stimuli.

#### Sub-populations

For each of the retinotopic and relational models (Fig. 3b), the top 20 units were selected based on cross-validated model accuracy. For these two resulting sub-populations, a smoothed firing rate was defined for a stimulus as the average of firing rates for all trials weighted by a Gaussian kernel in the (θ₀, θ₁) plane with width σ = 20°, evaluated with angular wrap-around to ensure there were no boundary effects.

#### Fisher information

We measure discriminability along a dimension in stimulus space as d′² = uᵀ J u, where J is the population’s 2 × 2 Fisher information matrix. Specifically, to measure d′² along a unit direction u in the (θ₀, θ₁) plane, we considered the two stimuli s_+_ = s + (Δ/2)·u and s_-_ = s - (Δ/2)·u, with Δ = 25°. For unit i, we compute

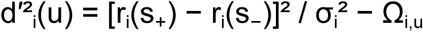

Where r_i_ is the firing rate of unit i, σ_i_² is the firing rate variance of unit i (pooled over the two stimuli s_+_ and s_-_), and Ω_i,s,u_ is a correction term for the fact that trials may count towards both stimuli due to smoothing:

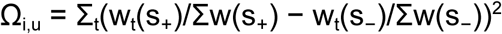

Where w_t_(s) is the Gaussian smoothing kernel centered at trial s and evaluated at trial t. We sum over units, so d′²(u) = Σ_i_d′²_i_(u).

Near the ∼80° separation floor, a kernel centered on a stimulus overlaps trials on one side only. Moving the stimulus across that edge changes the smoothed firing rates less than it would in the interior. This causes an underestimate of the discriminability along the direction perpendicular to that edge, hence why the vectors are parallel to the stimulus set boundaries.

#### Synthetic data

Panel b shows the same analysis as panel a except using firing rates of idealized neurons instead of recorded neurons. We defined these as:

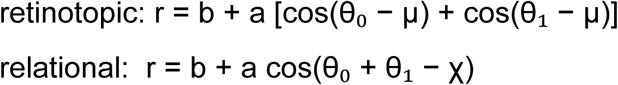

For a trial (θ_0_, θ_1_). We matched mean firing rates with the real population, so for each real neuron we had a corresponding synthetic neuron with b the mean firing rate of the real neuron. We let a = b/8 for retinotopic neurons and b/4 for relational neurons, so every neuron’s peak-to-trough modulation was 0.5. We let the preferred directions μ and χ evenly tile all angles, so the synthetic populations had no bias of preferred directions. Firing rates were Poisson sampled for synthetic neurons.

## ACKNOWLEDGEMENTS

We thank Rishi Rajalingham, Sujay Neupane, Alexandra Ferguson, and Hansem Sohn for their guidance in setting up the experimental rigs and developing our neurophysiology methods. We thank the CatalystNeuro team including Cody Baker, Luiz Tauffer, Felix Pei, Alessio Buccino, and Ben Dichter for their help open-sourcing our data. We thank Robert Desimone, Nancy Kanwisher, Gabriel Stine, and Amin Fakharian for their feedback on the manuscript.

## AUTHOR CONTRIBUTIONS

N.W., J.T., and M.J. designed the task. N.W. and J.G. collected the data. N.W. and M.J. developed the analyses. N.W. did the data processing and implemented the analyses. N.W., J.G., J.T., and M.J. wrote the manuscript.

## DATA AND CODE AVAILABILITY

The data used to generate all results is available in a public repository: https://dandiarchive.org/dandiset/000620

The code used to generate all results is available on a public repository: https://github.com/jazlab/relational_coding_2026

## SUPPLEMENTARY FIGURES

**Supplementary Figure 1.**
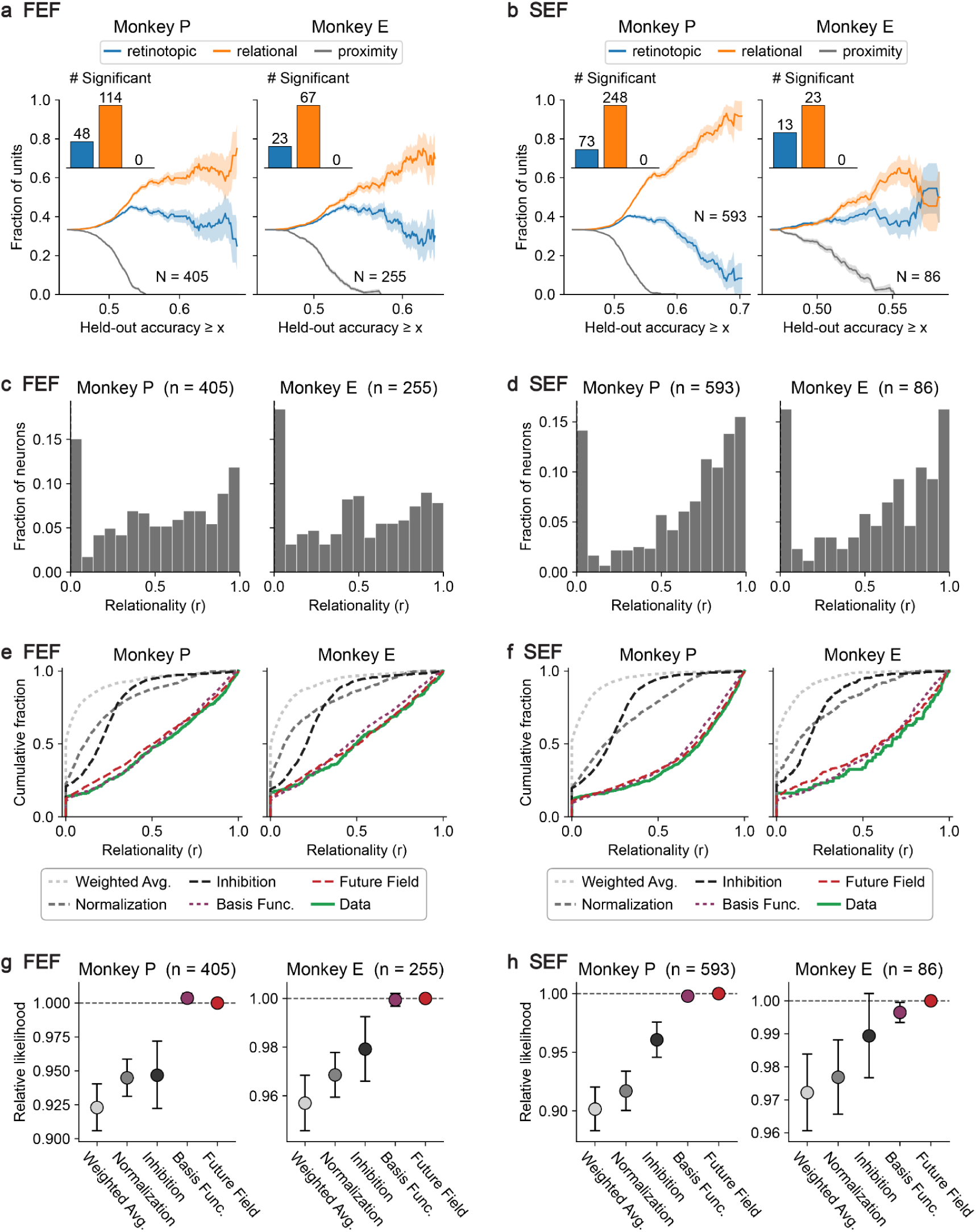
Results per brain area. **(a,c,e,g)** Same as main text Figure 3b, 4c, 4d, and 4g, except only considering FEF units. **(b,d,f,h)** Same as main text Figure 3b, 4c, 4d, and 4g, except only considering SEF units.

**Supplementary Figure 2.**
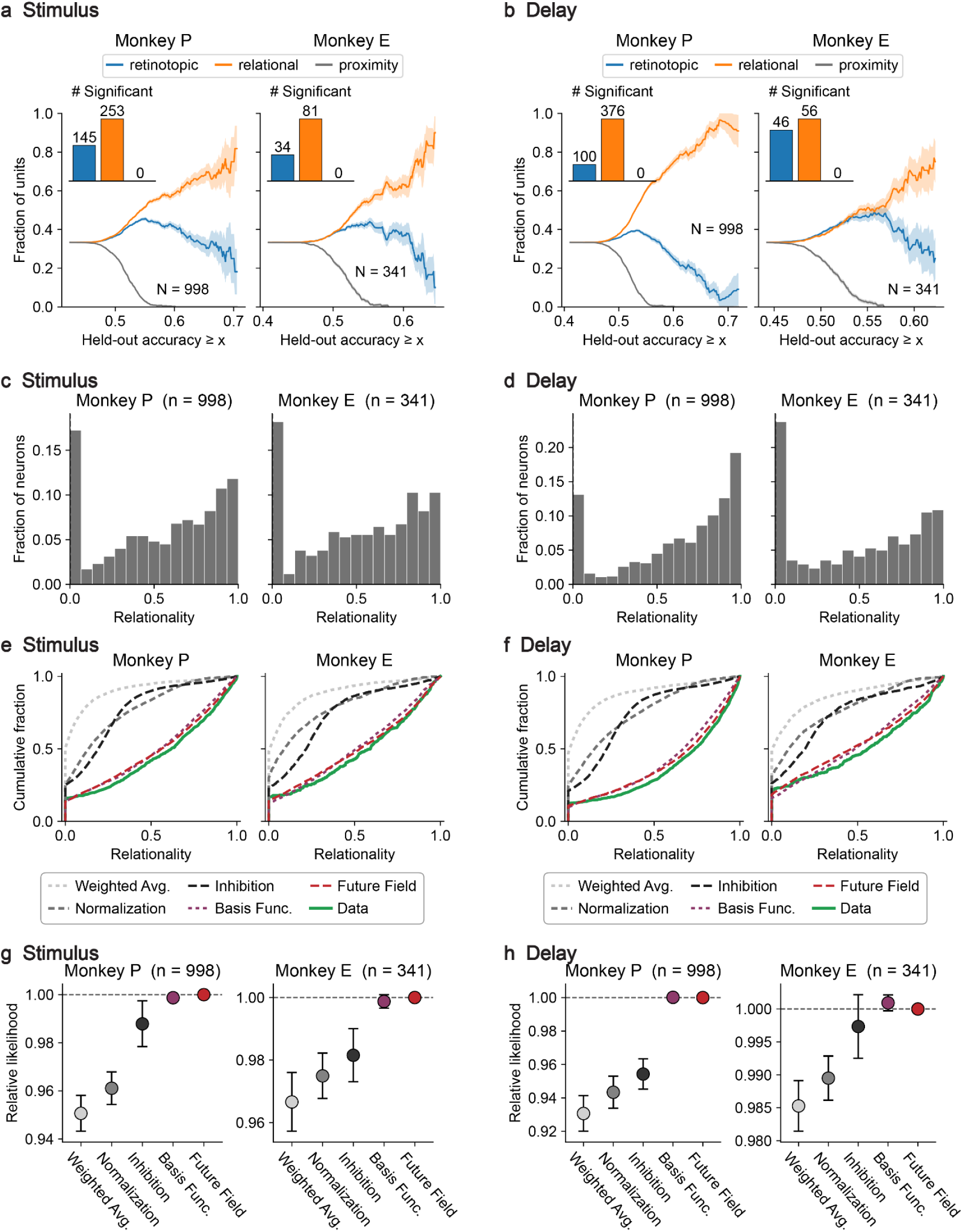
Results per task phase. **(a,c,e,g)** Same as main text Figure 3b, 4c, 4d, and 4g, except firing rates are computed only during the stimulus phase of the task. **(b,d,f,h)** Same as main text Figure 3b, 4c, 4d, and 4g, except firing rates are computed only during the delay phase of the task.

**Supplementary Figure 3.**
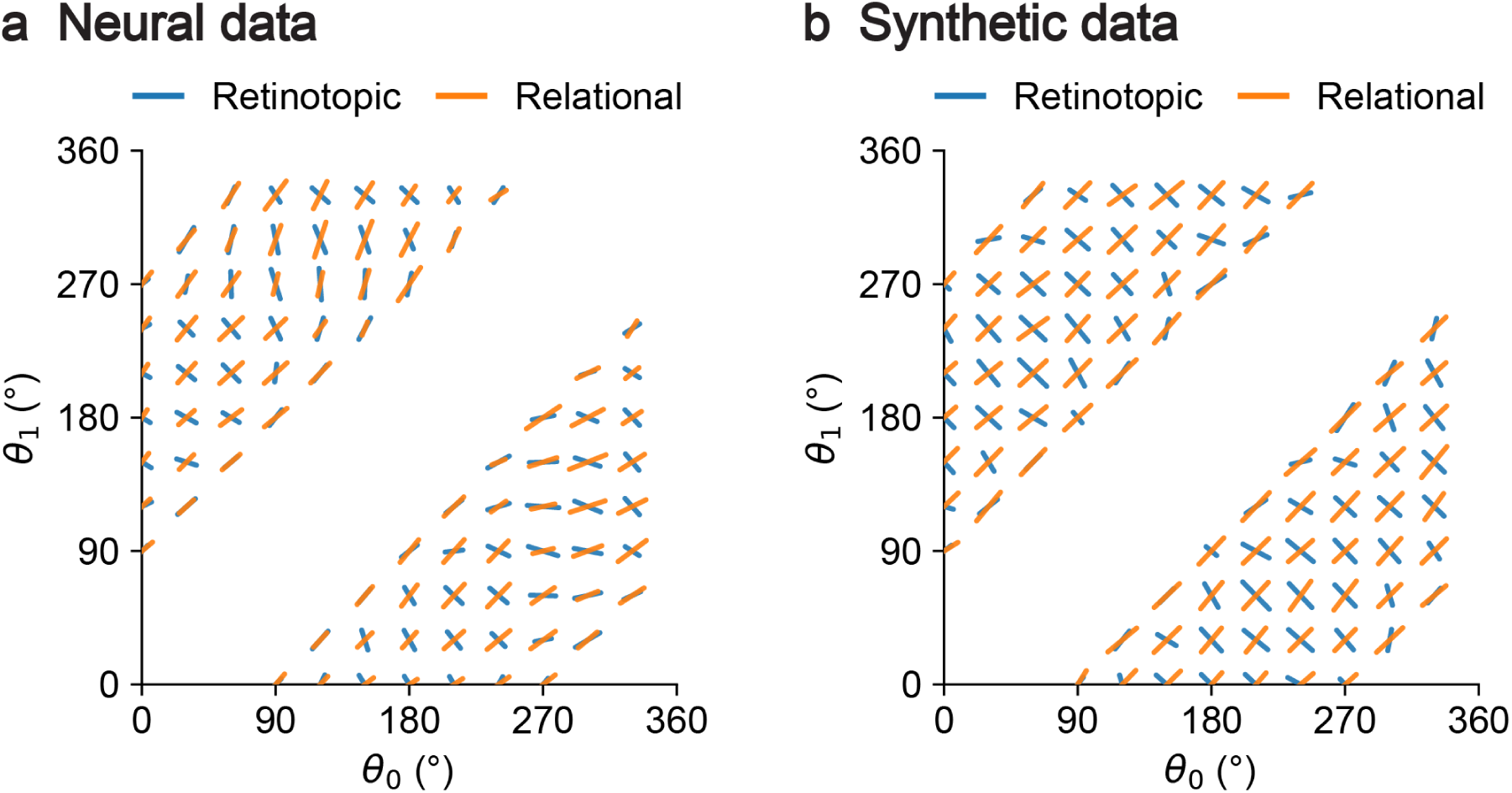
Dimensions of coding by relational and retinotopic units. **(a)** Neural data. Axes span the space of 2-object stimuli, as in Figure 1b (bottom). At each stimulus on a 30° grid, the blue segment shows the direction of maximum Fisher information of a retinotopic sub-population of units (the top 20 units by cross-validated retinotopic model accuracy, pooled across monkeys). Orange segments show the same but for a relational sub-population. The relational population segments have positive slope, indicating the relational population is best at distinguishing perturbations of both objects in the same direction, consistent with Figure 3c. The misalignment between blue and orange segments imply the retinotopic population provides complementary coding directions to the relational population. **(b)** Same as (a) except using synthetic data of idealized relational and retinotopic populations with preferred directions/orientations evenly tiling the circle.

**Supplementary Figure 4.**
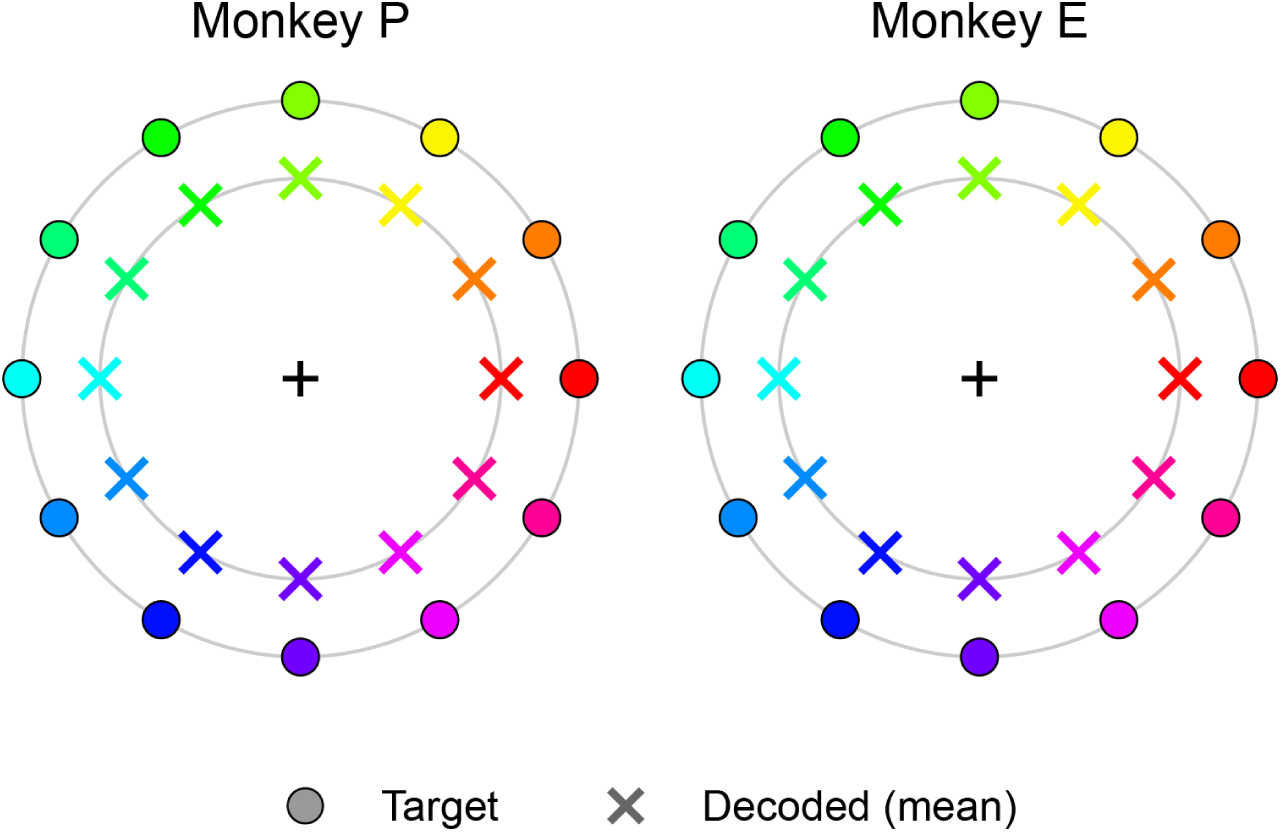
Response decoder on 1-object trials. The decoder from Fig. 3g/h was trained on 1-object conditions. Here are shown the decoder’s performance on those 1-object conditions. The exterior dots show 16 object angles, and the interior crosses show mean decoder output angle for these conditions. The standard deviation across trials was 1.83° for Monkey P and 1.85° for Monkey E.

**Supplementary Figure 5.**
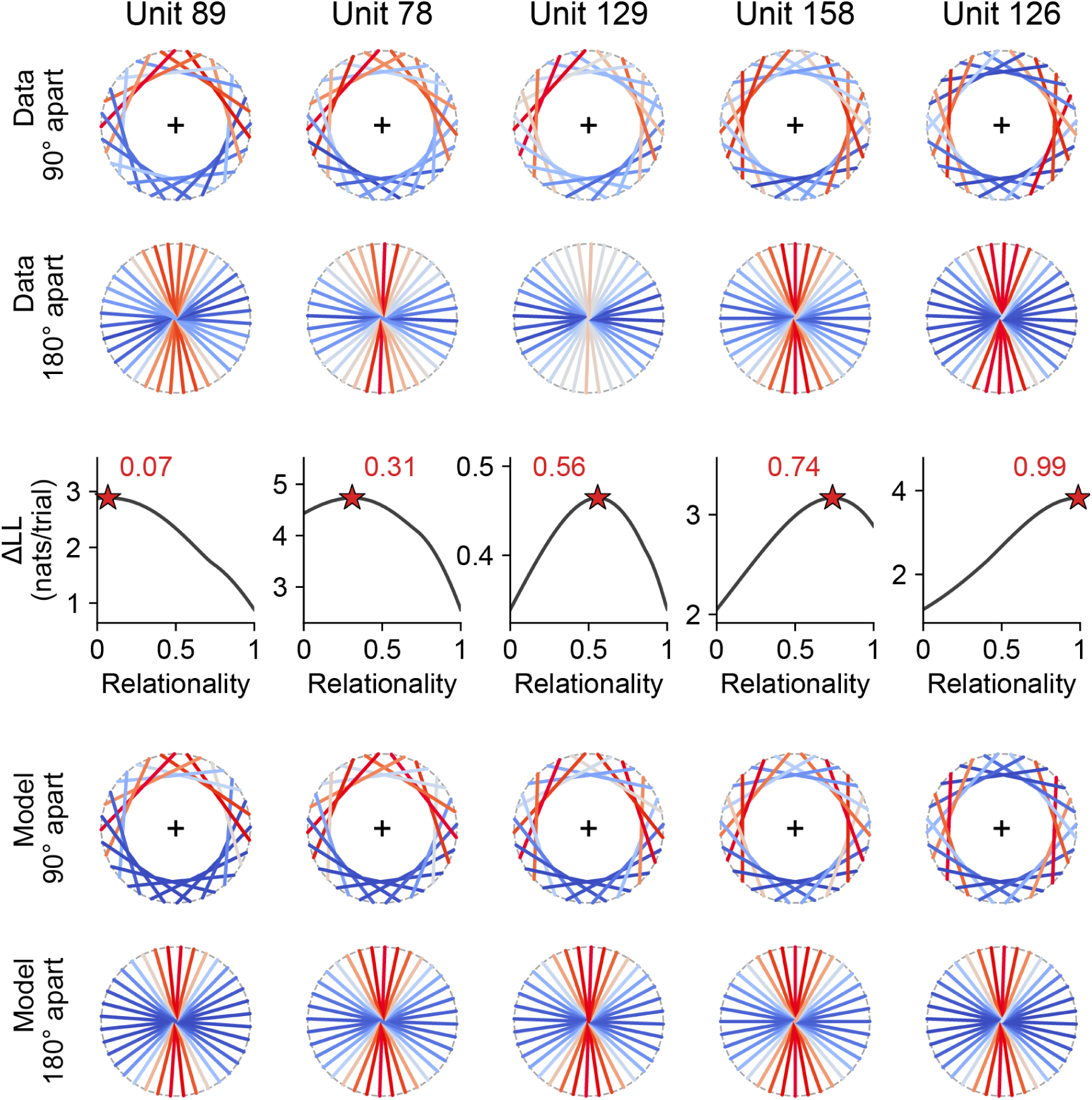
Relationality inference. Columns correspond to each unit in Figure 4a. Top two rows show tuning to 2-object conditions in the same format as Figure 2a-d, except rotated so that each unit’s 1-object receptive field is oriented upwards. Middle row shows relationality inference, where x-axis is relationality and y-axis is the corresponding likelihood of the data by fitting the relationality model with the given relationality. Stars show maxima, and labels report the inferred relationality value. Bottom two rows show the response tuning for 90° and 180° apart conditions for the relationality model (Methods).

**Supplementary Figure 6.**
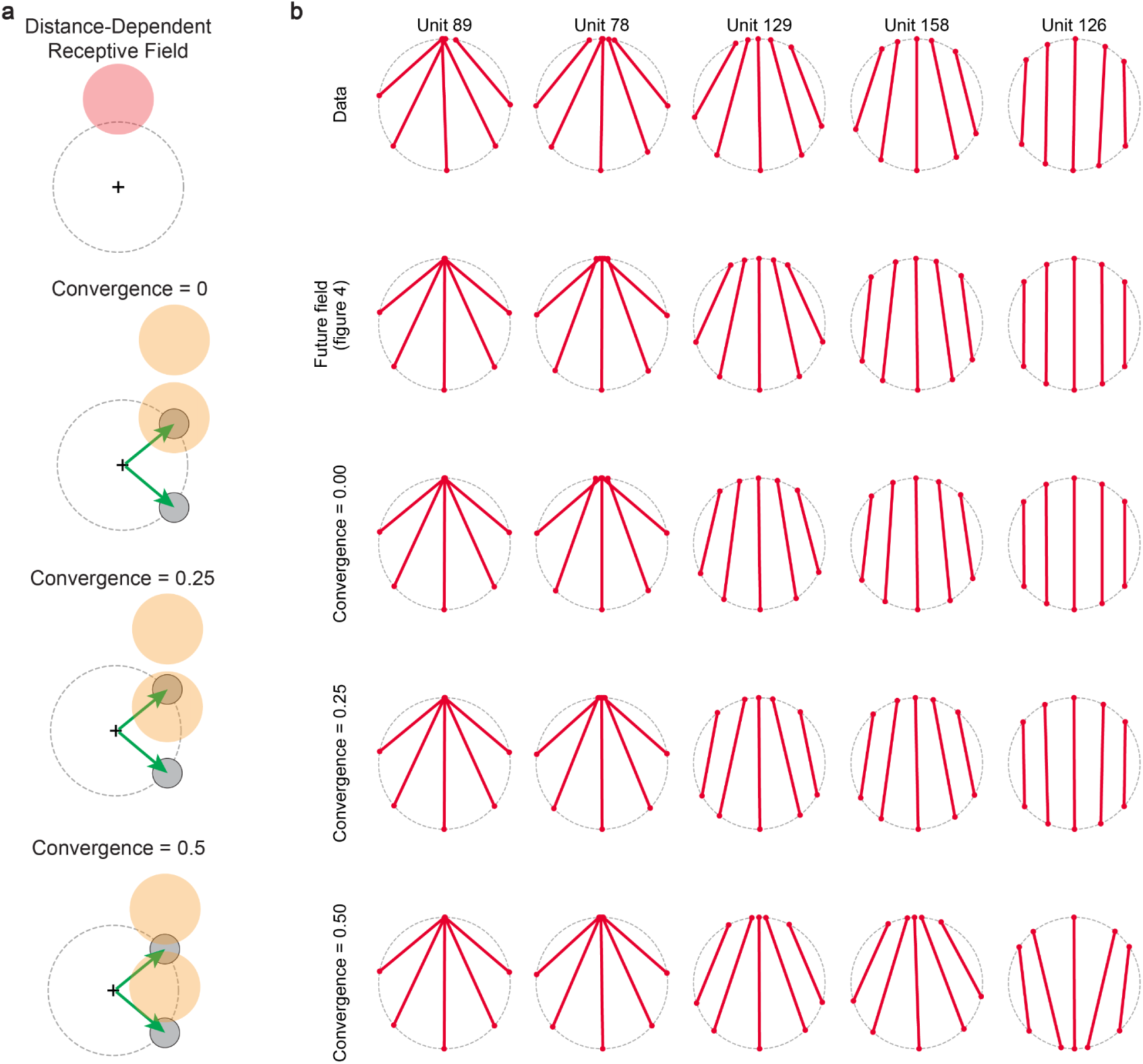
Convergent remapping. The future field model produces relational responses under convergent remapping. **(a)** Schematics of future fields under different degrees of convergence. For convergent remapping to be meaningful, receptive fields must depend on distance (top). The degree of convergence governs how close the future fields are to the saccade targets (bottom three schematics). **(b)** Considering the five units in Figure 4a (top row), their relationality was captured by the future field model with angular receptive fields (second row), and the future field model with Gaussian receptive fields with three degrees of convergence (bottom three rows). Each model was fitted to each neuron.

**Supplementary Figure 7.**
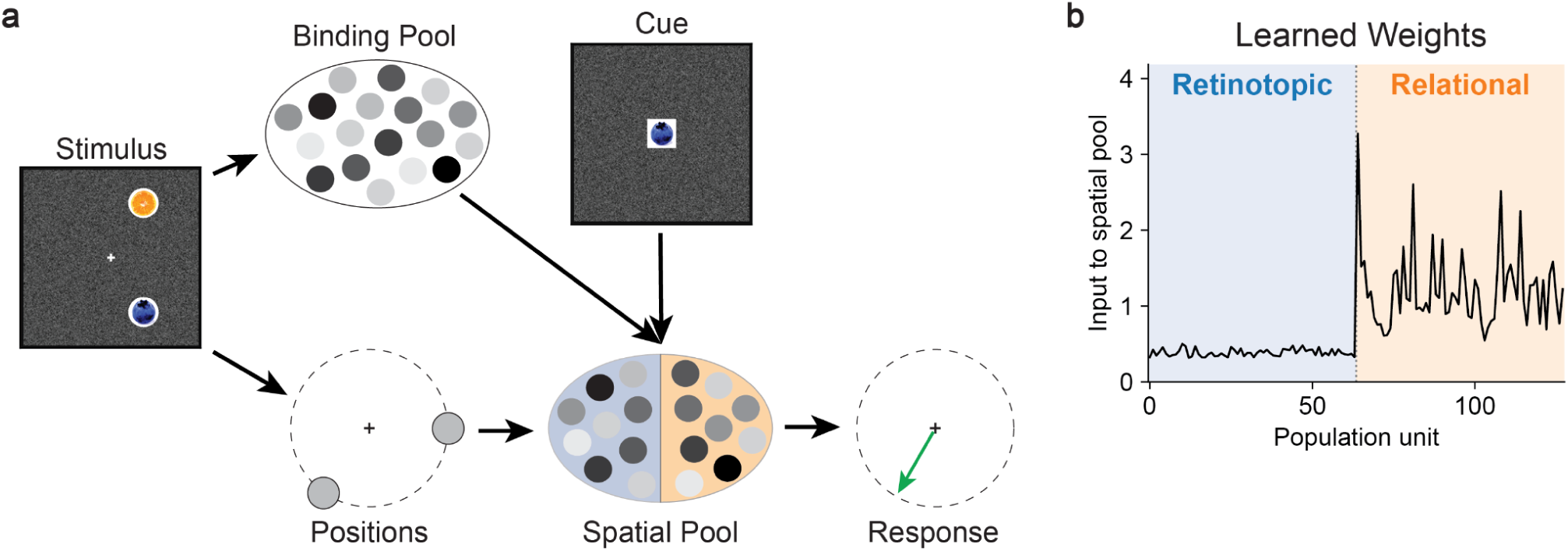
Binding model. **(a)** Model schematic. Spatial pool analogous to FEF and SEF consists of half idealized retinotopic units (blue shading) and half idealized relational units (orange shading). Binding pool receives stimulus object positions and identities, and projects with a cue to the spatial pool, which then generates a response saccade vector. **(b)** After training on the task, binding pool projects preferentially to relational units in the spatial pool. X-axis enumerates each of the model’s 128 units. Retinotopic units are on the left (blue shading) relational units are on the right (orange shading). Y-axis is the learned synaptic weight to each spatial pool unit, integrated over all binding pool units.

**Supplementary Figure 8.**
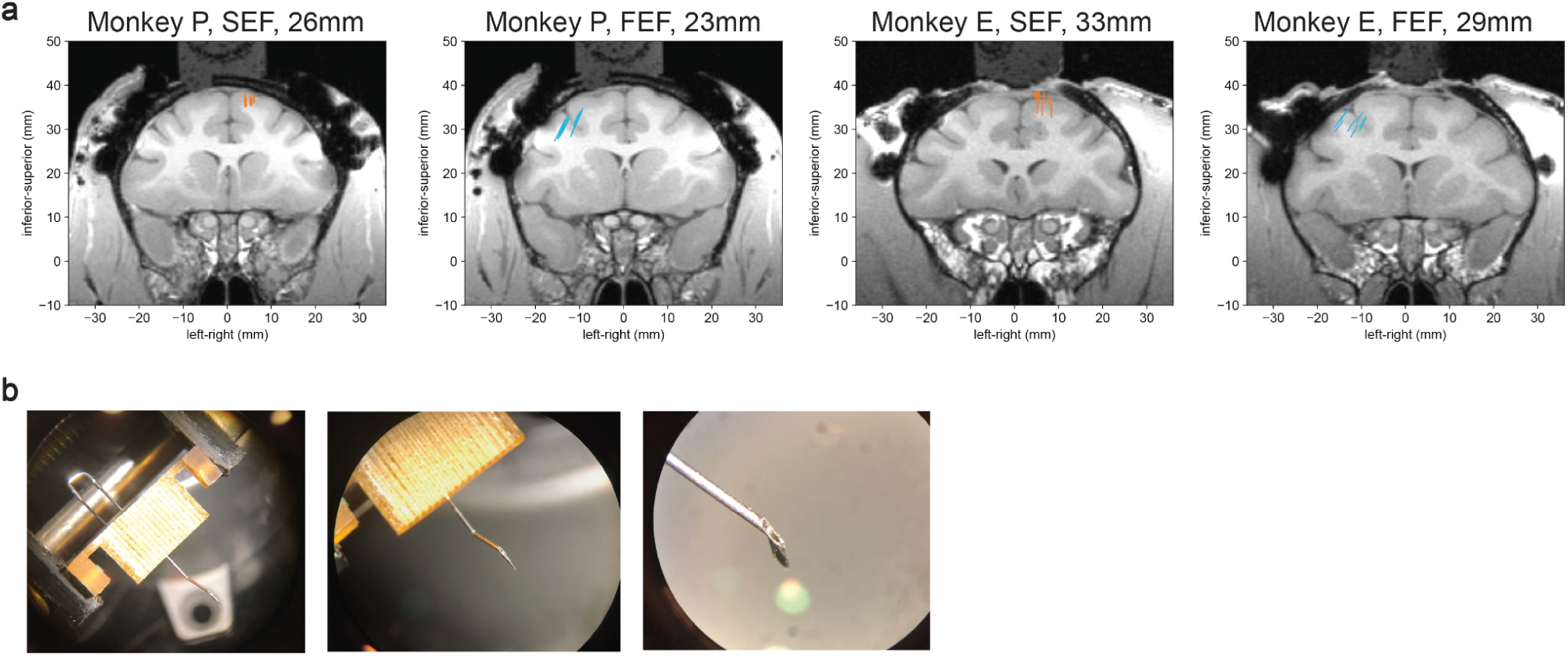
Recording targeting and guide tube system. **(a)** Coronal MRI slices showing probe trajectories for each recording area in each monkey. Millimeters in titles report MRI slices (in the Horsley-Clarke system relative to ear-bar zero). Each colored line is the trajectory of one electrode on one recording session, projected onto the MRI slice. Note that because of this projection, some trajectories were a few millimeters anterior or posterior of the shown slice. **(b)** Neuropixels recording guide tube, under three levels of magnification. The guide tube was made of a 23G syringe needle and hand-crafted with a Dremel and pliers. It passed through a grid and attached to a crimped J-shaped piece that then returned through the grid from the top to afford rotation stabilization. This was assembled and autoclaved before every session, then placed on the chamber for recording.

**Supplementary Figure 9.**
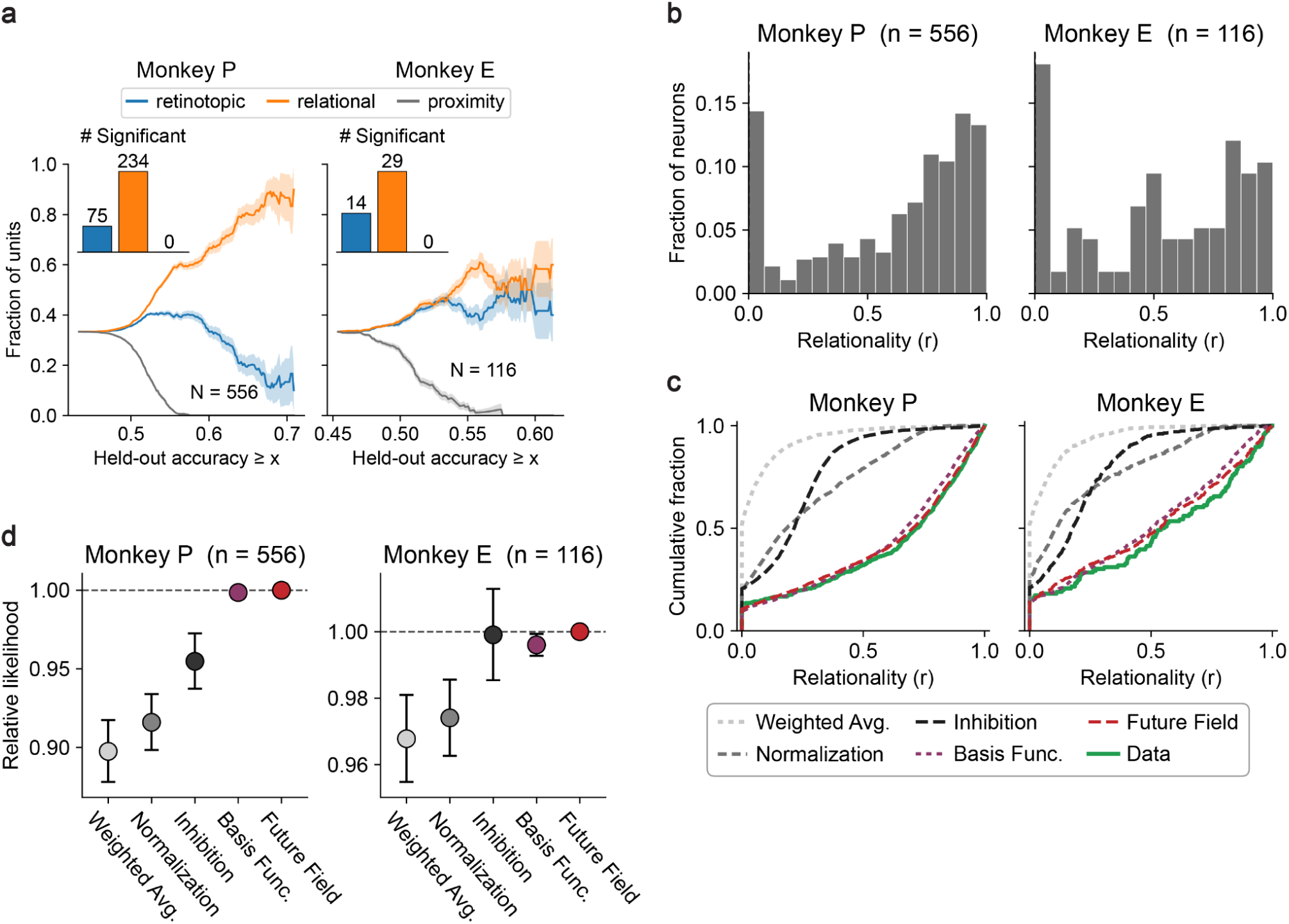
Results using single neurons only. Same as main text Figure 3b, 4c, 4d, 4g, except only using single neurons, excluding all units classified as multi-unit.

**Supplementary Figure 10.**
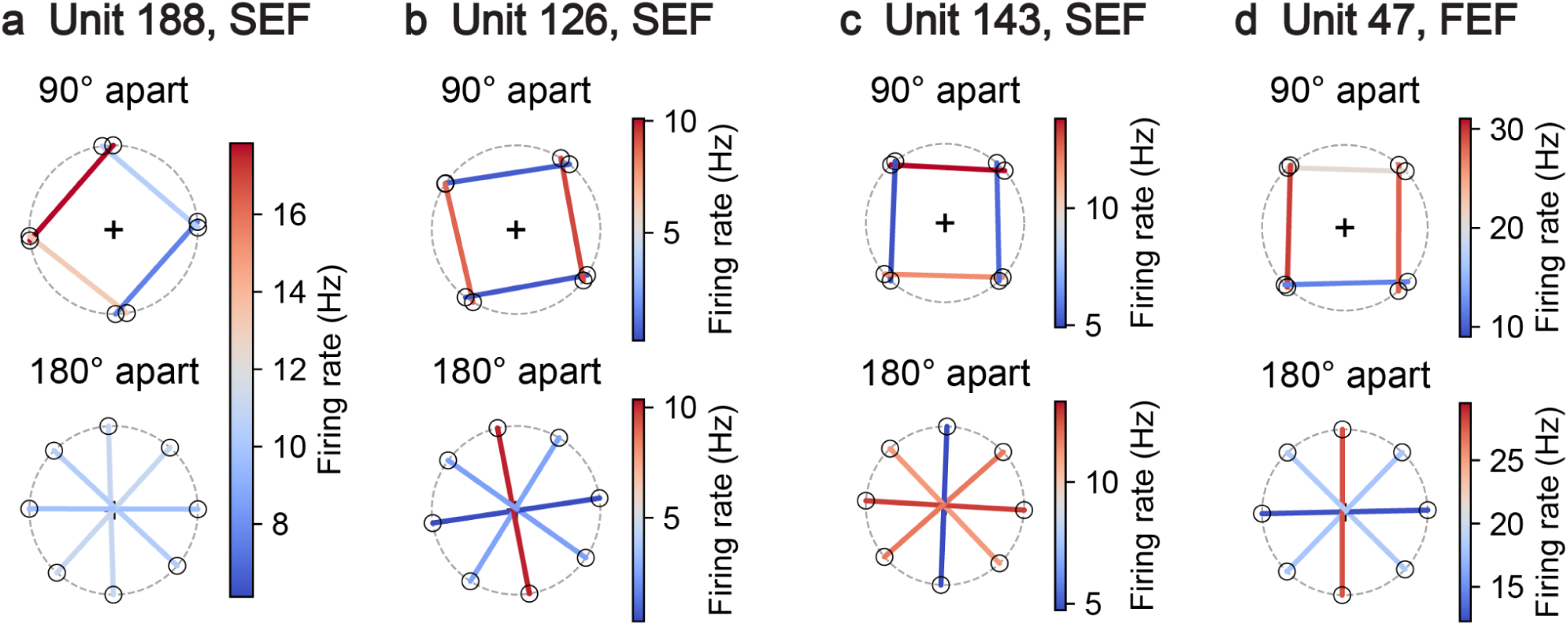
Empirical mean example stimuli. **(a-d)** Same stimulus plots as in Figure 2a-d, except every chord’s endpoints are the empirical means of the stimuli it averages over, not the prototype from which the average bin is computed.

